# A versatile, chemically-controlled DNA binding switch enables temporal modulation of Cas9-based effectors

**DOI:** 10.1101/2022.05.10.491425

**Authors:** Cindy T. Wei, Omri Peleg, Elhanan Borenstein, Dustin J. Maly, Douglas M. Fowler

## Abstract

CRISPR-Cas9’s RNA-guided genome targeting ability has been leveraged to develop a plethora of effectors including targeted transcriptional activators, DNA base editors, and DNA prime editors. Although systems for inducibly modulating Cas9 activity have been developed, current approaches for conferring temporal control require extensive screening of functional protein components. A simpler and general strategy for conferring temporal control over diverse Cas9-based effector activities is needed. Here we describe a versatile chemically-controlled and rapidly-activated DNA binding Cas9 module (ciCas9) that is able to confer temporal control over a variety of Cas9-based effectors. Using the ciCas9 module, we engineer temporally-controlled cytidine and adenine DNA base editors. We employ the ciCas9 base editors to reveal that *in vivo* bystander editing kinetics occurs via a dependent process where editing at a preferred nucleotide position increases the frequency of edits at a second nucleotide within a target site. Finally, we demonstrate the versatility of the ciCas9 module by creating a temporally-controlled transcriptional activator, a dual cytidine and adenine base editor, and a prime editor.

## Introduction

The CRISPR-Cas9 system consists of a Cas9 endonuclease that can be targeted to any location within a genome using a single guide RNA (sgRNA) encoding a 20 nucleotide targeting sequence^1–3^. The CRISPR-Cas9 system is commonly used to create genomic double-strand breaks (DSBs) to facilitate incorporation of desired DNA edits at specific loci via homology directed repair (HDR) or to generate indels to knock out specific genetic elements via non-homologous end joining (NHEJ)^4^. Engineered Cas9-based effectors have enabled a plethora of applications beyond DSB generation for DNA editing^4^. For example, catalytically-inactive Cas9 (dCas9) has been fused to transcriptional activators or repressors to modulate gene expression and to chromatin modifiers for targeted epigenome editing^5^. Nickase Cas9 (nCas9) has been fused to DNA deaminase enzymes to yield cytidine to thymidine and adenine to guanidine DNA base editors^6, 7^. Dual C-to-T and A-to-G base editors and C-to-G base editors have also been engineered^8–12^. Recently, prime editing has been developed to introduce precise DNA edits using an RNA template^13^. Thus, the CRISPR-Cas9 system has proven to be extraordinarily versatile.

Engineered inducible Cas9 variants have also been developed to provide temporal control over targeted DSB generation and subsequent DNA editing^14–26^. Such temporally-controlled Cas9s have been used in a variety of applications including studying the kinetics of CRISPR-Cas9 DNA editing and the kinetics of DNA repair after a DSB, as well as to engineer systems that record biological events in cells^26–29^. Beyond temporally-controlled Cas9s for generating DSBs, temporally-controlled versions of other Cas9-based effectors have also been engineered. Examples include temporally-controlled dCas9-based DNA transcription and chromatin modifiers capable of turning on or off gene expression^24, 30–33^ and split-engineered base editors (seBEs) that allow for temporal control of C-to-T base editing^34^.

Although temporal control of some Cas9-based effectors has been achieved, existing systems comprise a patchwork of approaches that do not cover all important Cas9-based effector activities (Supplemental Table 1). Moreover, these existing systems are complicated by the fact that they often require screening of split enzymes to confer temporal control over specific Cas9-based effector domains. These approaches can be laborious and are not easily applicable to all effectors. A generalizable system to engineer temporal control of all Cas9-based effectors would be based on a single component, would be rapidly activatable, and would allow precise tuning of activity. Since RNA-guided Cas9 binding to DNA is common to all Cas9-based effectors, we hypothesized that control of DNA binding activity would enable engineering of all Cas9-based effector systems.

We previously developed a single component, temporally-controlled Cas9 protein, ciCas9, that contains a tightly autoinhibited switch that can be rapidly activated with a potent small molecule^26^. Here, we show that the ciCas9 switch can serve as a general platform for conferring temporal control over a wide range of Cas9-based effectors. We develop a chemically-controlled transcription factor, dciCas9-VPR, and use it to show that the ciCas9 switch functions by governing DNA target site binding. We then use the ciCas9 switch to engineer chemically-controlled base editors, allowing robust temporal control over C-to-T and A-to-G DNA editing. We employ these chemically-controlled base editors to explore, for the first time, how nucleotide position within a target site and early base editing kinetics affect editing outcomes. We also dissect the kinetics of allele formation, elucidating the order in which nucleotides are edited and revealing how base editing at one nucleotide in the target site influences bystander edits at other nucleotides within the same target. Finally, we highlight the versatility of the ciCas9 switch by engineering chemically-controlled dual A-to-G and C-to-T base editors and DNA prime editors whose activity can be controlled with high temporal precision.

## Results

### The ciCas9 switch facilitates chemically-controlled DNA target site binding

We previously developed a chemically-controlled Cas9 variant (ciCas9) in which the REC2 domain was replaced with Bcl-xL and to which a BH3 peptide was appended^26, 35, 36^. Bcl-xL and BH3 form a tight intramolecular complex that inhibits Cas9 activity (Fig. 1a). In the basal state, autoinhibited ciCas9 possesses low activity, but addition of a small molecule (A-1155463, hereafter A115) disrupts the interaction between Bcl-xL and the BH3 peptide resulting in dose-dependent generation of double-stranded breaks (DSBs) at target sites within minutes^26^. We reasoned that the single-protein architecture and rapid activation kinetics of ciCas9 could serve as a versatile platform for conferring chemical control over diverse Cas9 effector activities. Successful application of the ciCas9 switch to Cas9-based effectors requires that the switch modulates DNA target site binding as opposed to another mechanism such as altering Cas9 enzymatic activity (Fig. 1a). To test whether ciCas9’s autoinhibitory switch controls DNA target site binding *in vivo*, we measured transcriptional activation, which relies on Cas9 localization through DNA binding rather than Cas9 nuclease activity. Thus, we fused the transcriptional activator Vp64-p65-Rta (VPR) to the C-terminus of catalytically dead ciCas9 (dciCas9) and tested its ability to promote expression of CXCR4 (Fig. 1b, c). A115 treatment of HEK-293T cells expressing dciCas9-VPR resulted in induction of CXCR4 expression, supporting the DNA-blocking autoinhibition mechanism of dciCas9. Consistent with ciCas9 acting as a chemically-controlled DNA target site binding switch, we also found that unmodified dciCas9 functions in a multi-protein component transcriptional activation assay with previously reported scaffold RNAs (scRNA) (Supplemental Fig. 1)^37^. Therefore, ciCas9 is unable to bind DNA target sites in its autoinhibited state, and release of autoinhibition by A115 addition allows ciCas9 to bind. Finally, we demonstrated that dciCas9-VPR transcriptional activation could be dose-dependently tuned by targeting dciCas9-VPR to a synthetic EGFP reporter in the presence of different amounts of A115 (Fig. 1d; Supplemental Fig. 2). Thus, the ciCas9 switch modulates DNA binding, a process common to all Cas9-based effectors, and can be temporally and dose-dependently controlled with small molecules.

**Figure 1.**
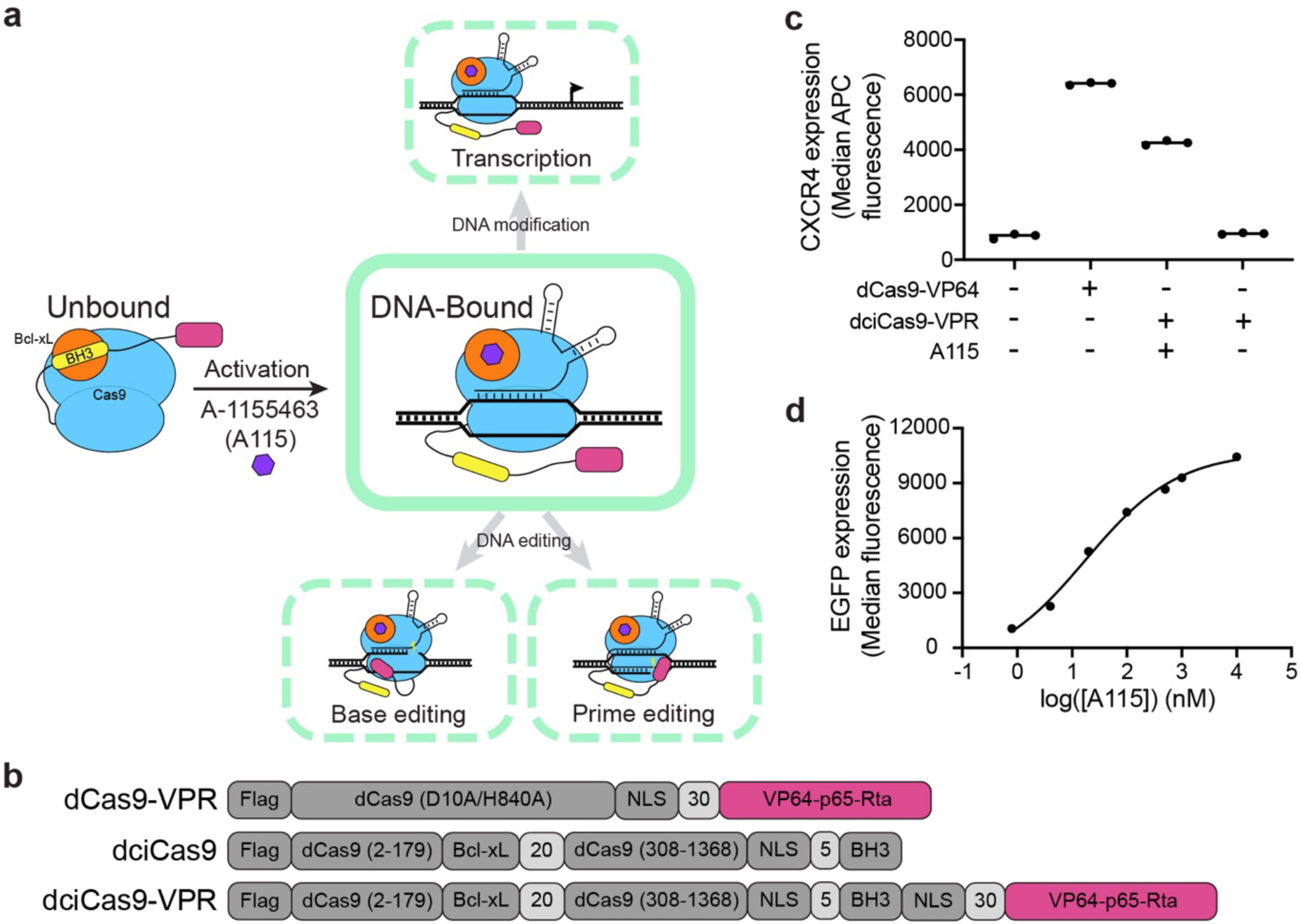
The ciCas9 switch can be used as a framework to create chemically-controlled Cas9-based effectors. **a)** Schematic showing how the ciCas9 switch can be used to engineer different chemically-controlled Cas9-based effectors. **b)** Domain schematic of catalytically-dead Cas9 (dCas9) and ciCas9 (dciCas9) fused to the transcriptional activator VP64-p65-Rta (VPR). **c)** Activation of CXCR4 expression with dCas9-VPR or dciCas9-VPR targeted to the promoter region in HEK-293T cells in the presence or absence of 1 μM A115. Cells were stained with a fluorescently labeled anti-CXCR4 antibody. Three cell culture replicates are shown, with a line indicating the mean. **d)** Activation of an EGFP reporter locus downstream of an EMX1 target sequence (EMX1-EGFP) using dciCas9-VPR and a range of A115 doses added to HEK-293 TREx FlpIn cells. Cells were treated with A115 for 48 hr prior to flow cytometry analysis. Points represent the mean of median EGFP fluorescence ± SEM of three cell culture replicates. Line shows a non-linear fit of log(agonist) vs. response - variable slope calculation in GraphPad Prism 9.

**Figure 2.**
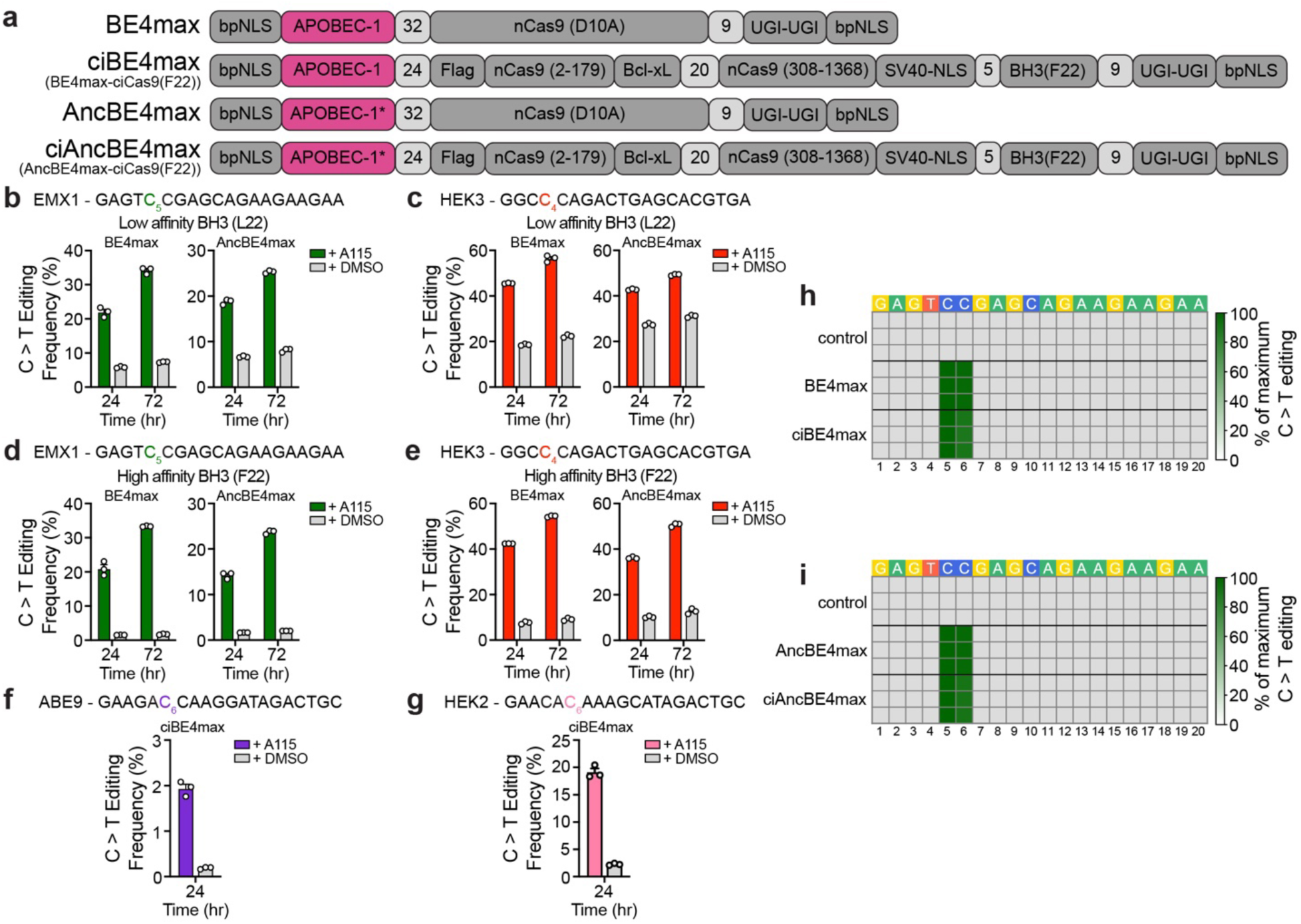
The ciCas9 switch can be used to create chemically-controlled cytidine base editors. **a)** Domain arrangements of the unmodified BE4max and AncBE4max base editors and of the optimized chemically-controlled base editors ciBE4max and ciAncBE4max. **b, c)** C-to-T editing frequencies of chemically-controlled BE4max and AncBE4max constructs containing the L22 BH3 peptide variant at the EMX1 **(b)** and HEK3 **(c)** target sites in HEK-293T cells treated with A115 or DMSO for 24 and 72 hr. **d, e)** C-to-T editing frequencies of chemically-controlled BE4max and AncBE4max constructs containing the F22 BH3 peptide variant (ciBE4max and ciAncBE4max) at the EMX1 **(d)** and HEK3 **(e)** target sites in HEK-293T cells treated with A115 or DMSO for 24 and 72 hr. **f, g)** C-to-T editing frequencies of ciBE4max and ciAncBE4max at the ABE9 **(f)** and HEK2 **(g)** target sites in HEK-293T cells treated with A115 or DMSO for 24 and 72 hr. In **(b-g)** editing was quantified at the single nucleotide within a target site that is colored in the target sequence. Bars show mean editing frequency ± SEM of three cell culture replicates, with white circles representing individual replicates. **h-i)** Heatmaps of BE4max, ciBE4max **(h)** and AncBE4max, ciAncBE4max **(i)** editing as a percentage of the highest edited nucleotide for each editor throughout the entire EMX1 target site. Each row shows an individual cell culture replicate. BE4max and AncBE4max editing frequencies were quantified at 72 hr after transfection and ciBE4max and ciAncBE4max editing frequencies were quantified at 72 hr after 1 μM A115 addition to HEK-293T cells. The control shows untransfected cells harvested at the same time as ciBE4max and ciAncBE4max. The numbers below the heatmaps show the position of the nucleotide from the most PAM-distal nucleotide.

### The ciCas9 switch can be used to create chemically-controlled DNA base editors

To further explore the utility of the ciCas9 switch, we created chemically-controlled cytidine base editors by fusing the BE4max or AncBE4max deaminases to a ciCas9 nickase (nciCas9), preserving the original domain arrangements (Fig. 2a)^38^. We then transfected the chemically-controlled cytidine base editors into HEK-293T cells and determined background base editing (DMSO treatment) and maximum base editing when the ciCas9 switch was fully activated with a high concentration of A115 (1 μM) using next-generation sequencing (Fig. 2b, c; Supplemental Fig. 3). For both chemically-controlled cytidine base editors, we observed modest DMSO background editing with robust A115-activated editing after 24 and 72 hr of activation with A115. The HEK3 target site accumulated more background edits than the EMX1 target site. In an attempt to maximize overall editing and reduce background, we modified the epitope tag, nuclear localization sequence, peptide linker lengths, and codon optimization of the chemically-controlled cytidine base editors (Supplemental Figs. 3-5). However, these factors did not have an appreciable impact on chemically-controlled editing.

**Figure 3.**
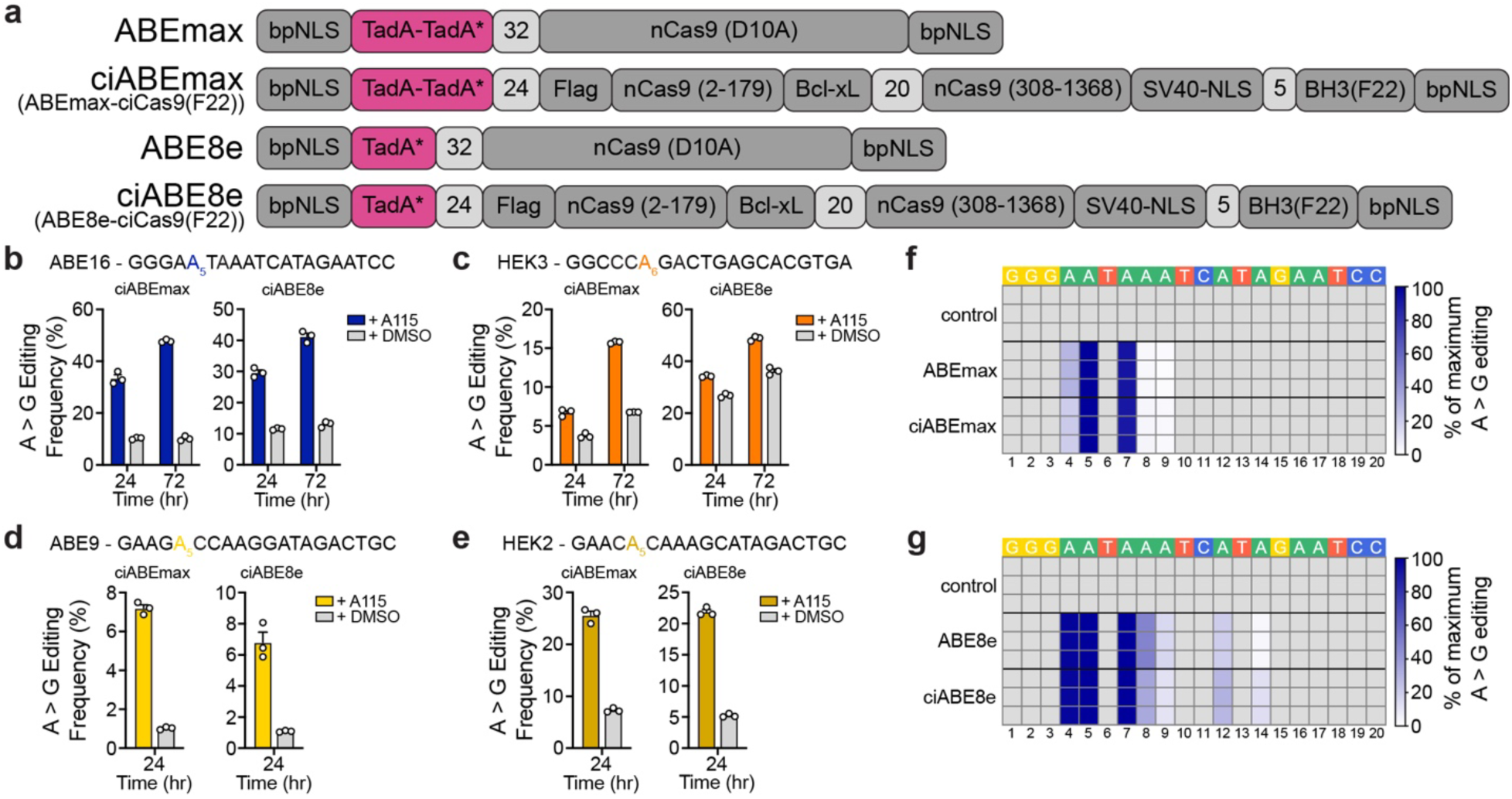
The ciCas9 switch can be used to create chemically-controlled adenine base editors. **a)** Domain arrangements of the unmodified ABEmax and ABE8e base editors and the optimized chemically-controlled base editors ciABEmax and ciABE8e. **b-e)** A-to-G editing frequencies of ciABEmax and ciABE8e base editors at the ABE16 **(b)**, HEK3 **(c)**, ABE9 **(d)**, and HEK2 **(e)** target sites in HEK-293T cells treated with A115 or DMSO for the times indicated. Editing is quantified at a single nucleotide within a target site that is colored in the target sequence. Bars show mean editing frequency ± SEM of three cell culture replicates, with white circles showing individual replicates. **f, g)** Heatmaps of ABEmax, ciABEmax **(f)** and ABE8e, ciABE8e **(g)** editing as a percentage of the highest edited nucleotide for each editor throughout the entire ABE16 target site. Each row shows an individual cell culture replicate. ABEmax and ABE8e editing frequencies were quantified at 72 hr after transfection and ciABEmax and ciABE8e editing frequencies were quantified at 72 hr after 1 μM A115 addition to HEK-293T cells. The control shows untransfected cells harvested at the same time as ciABEmax and ciABE8e. The numbers below the heatmaps show the position of the nucleotide from the most PAM-distal nucleotide.

**Figure 5.**
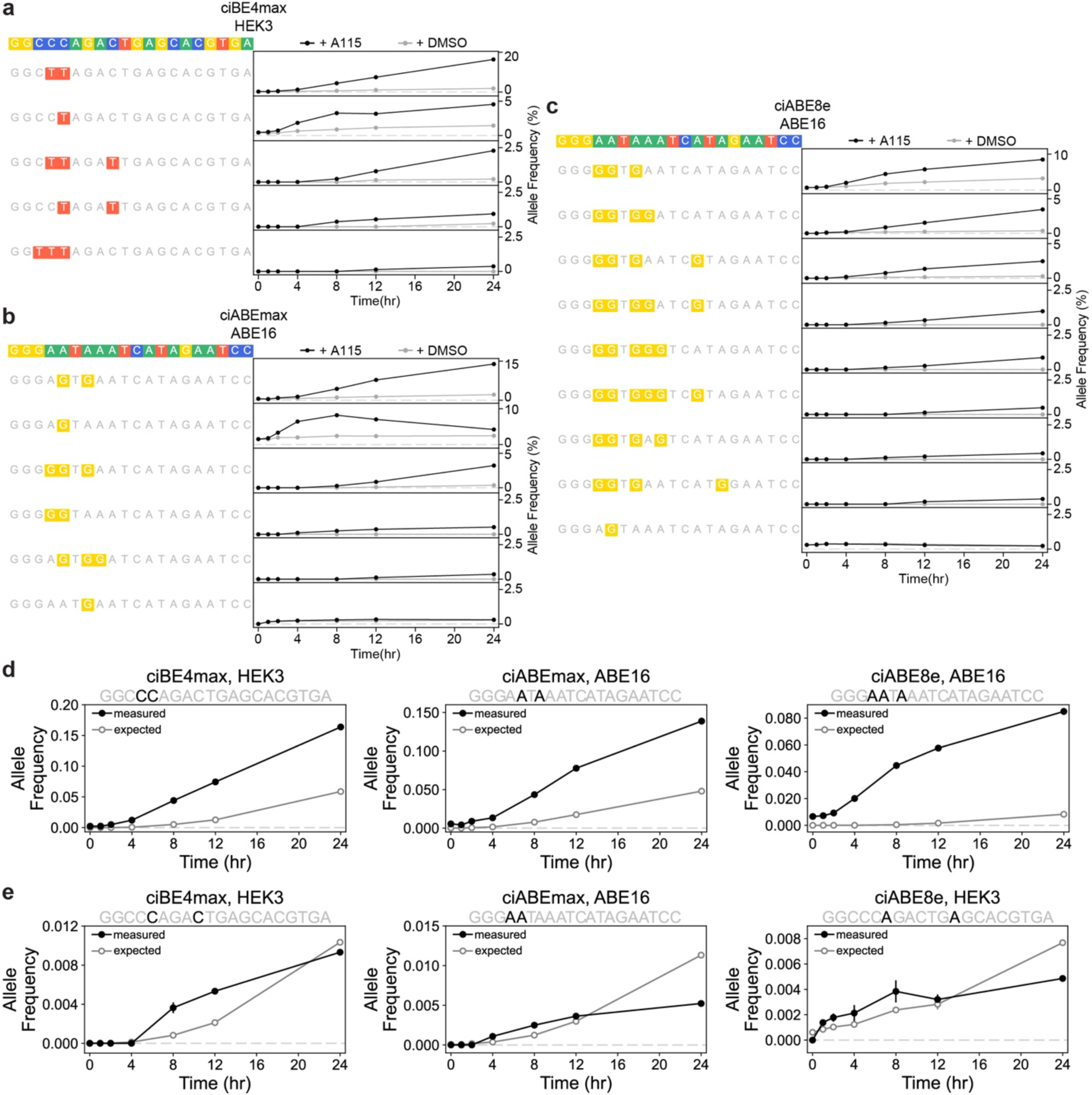
Chemically-controlled base editors reveal the kinetics of multiply-edited allele formation and nucleotide editing dependency. **a-c)** Time course of allele formation by ciBE4max at the HEK3 target site **(a)**, ciABEmax at the ABE16 target site **(c)**, and ciABE8e at the ABE16 target site **(c)**in HEK-293T cells treated with 1 μM A115 or DMSO. Cells were harvested and editing was quantified at specified time points after A115 addition. Black lines and circles show editing with 1 μM A115, gray lines and circles show editing with DMSO. Data represented as mean allele frequency ± SEM of 3 cell culture replicates. **d)** Examples of measured (black with solid circles) and expected (gray with open circles) allele frequencies over time by ciBE4max (left), ciABEmax (center), ciABE8e (right) that show a dependent model of base editing for multiply-edited alleles. **e)** Examples of measured (black with solid circles) and expected (gray with open circles) allele frequencies over time created by ciBE4max (left), ciABEmax (center), ciABE8e (right) that show an ambiguous model of base editing for multiply-edited alleles. Measured data represented as mean editing frequency ± SEM of 3 cell culture replicates. Expected editing frequency represented as mean expected editing frequency ± relative error. Calculations for expected frequency and relative error described in Materials and Methods.

We reasoned that background editing was due to the nciCas9 switch not being sufficiently closed, with transient dissociations of the BH3/Bcl-xL complex allowing DNA binding and subsequent base editing. Thus, to minimize background, we tested a higher affinity BH3 peptide variant, F22, that provides greater autoinhibition of nciCas9 activity (Figs. 2d, e)^35^. The higher affinity F22 variant did not appreciably reduce editing at 24 or 72 hr after A115 addition compared to the lower affinity L22 variant (Figs. 2b-e). However, the F22 variants of both chemically-controlled cytidine base editors demonstrated lower DMSO background editing at both the EMX1 and HEK3 target sites. To verify F22 variant performance, we evaluated chemically-controlled BE4max at the ABE9 and HEK2 target sites, where we observed similarly low background (Figs. 2f, g). Thus, the dynamic range of chemically-controlled cytidine base editors can be increased by strengthening the autoinhibitory interaction between Bcl-xL and the BH3 peptide (Supplemental Figs. 4c, 4e, 5c, 5e). We used the F22 variants of both cytidine base editors, which we hereafter refer to as ciBE4max and ciAncBE4max, for all subsequent studies.

**Figure 4.**
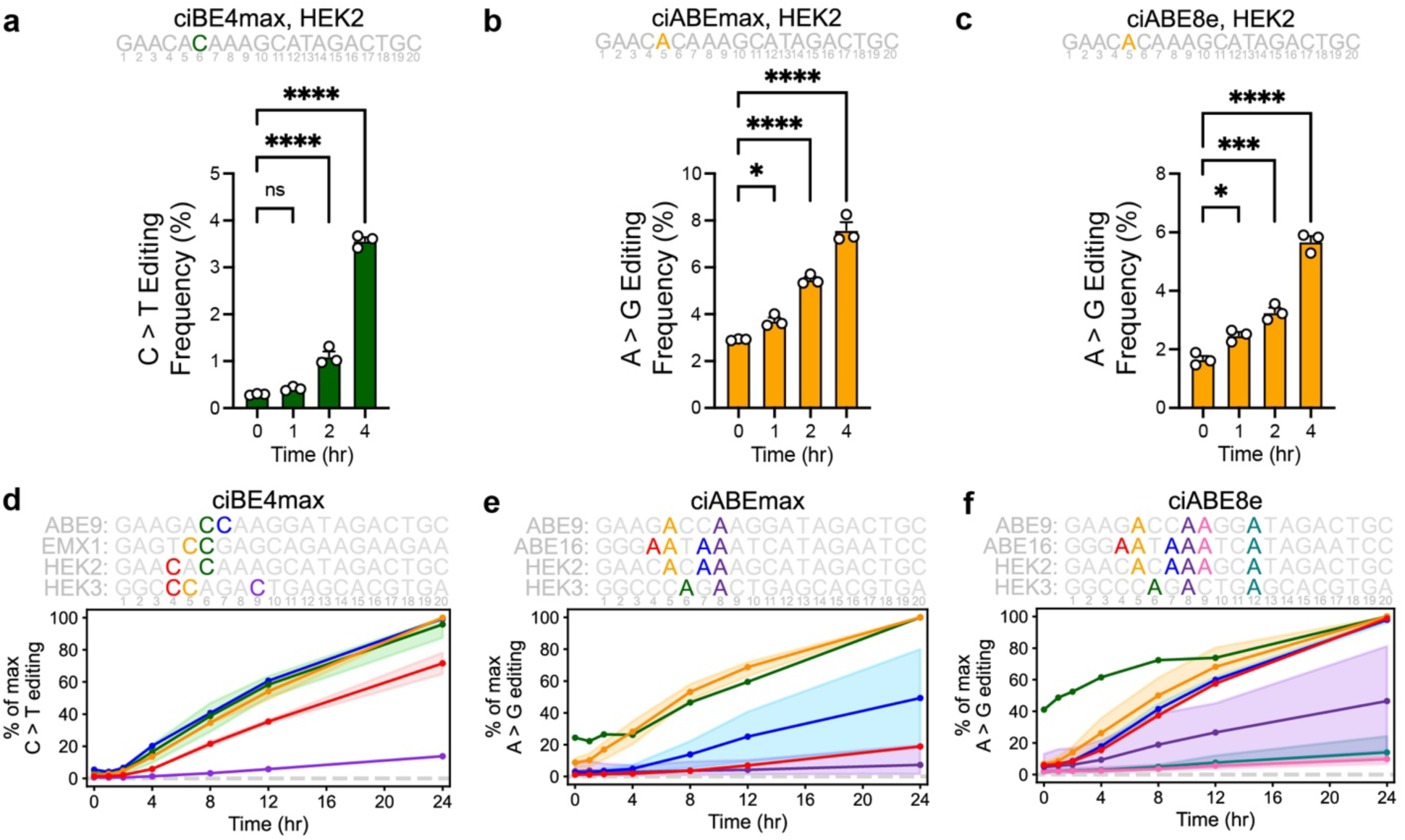
Chemically-controlled base editors reveal the effect of nucleotide position on editing kinetics. **a-c)** Early editing time courses at the HEK2 target site with the ciBE4max **(a)**, ciABEmax **(b)**, and ciABE8e **(c)** base editors in HEK-293T cells treated with 1 μM A115. Editing frequencies are for the nucleotide colored in the HEK2 target sequence. Numbers underneath the target sequence show the position of the nucleotide from the most PAM-distal nucleotide. Bars show mean editing ± SEM of 3 cell culture replicates with white circles representing individual replicates. Significance of editing at different time points were compared to editing frequency at 0 hr using a one-way ANOVA, statistical values shown in Supplemental Table 2. **d-f)** Normalized editing time courses for ciBE4max **(d)**, ciABEmax **(e)**, and ciABE8e **(f)** in HEK-293T cells treated with 1 μM A115 . Time courses were normalized within each target sequence where the highest edited nucleotide within each target site at 24 hr after A115 addition was set to 100%. Lines show the mean normalized editing frequency at that position of all target sites listed, and shading shows the range between maximum and minimum normalized editing frequency at that position across all target sites. Color corresponds to the nucleotide positions within each target sequence, shown above each plot. Numbers underneath the target sequences show the position of the nucleotide from the most PAM-distal nucleotide.

Having engineered a set of robust, chemically-controlled cytidine base editors, we validated that they edit DNA similarly to their parent base editors by exploring their key properties. Both chemically-controlled cytidine base editors were only able to edit the intended on-target site in the presence of an sgRNA (Supplemental Figs. 9a-d). Furthermore, both edited the same nucleotides within a target site to a similar degree as the parental versions (Figs. 2h-i; Supplemental Figs. 10a-b), with minimal indel formation at the target site (Supplemental Figs. 11a-d). Finally, off-target DNA base editing occured at similar or lower magnitudes and at the same nucleotide positions compared to the parental base editors at all off-target sites investigated (Supplemental Figs. 12a-b). Thus, our chemically-controlled cytidine base editors do not appear to appreciably impact R-loop formation, positioning and dynamics of the DNA deaminase enzymes, or unwanted off-target DNA base editing activities relative to the parental versions. Thus, nciCas9 can be used as a direct replacement of nCas9, and simply appending the deaminase components results in chemical control of base editing.

We next engineered chemically-controlled adenine base editors by fusing either the ABEmax or ABE8e deaminases to the nciCas9 switch in the same domain arrangements as the unmodified ABEmax and ABE8e base editors (Fig. 3a)^38, 39^. We observed robust editing for both chemically-controlled adenine base editors when fully activated (Figs. 3b-e). Similar to the chemically-controlled cytidine base editors, the higher affinity F22 BH3 variant was able to improve the dynamic range of base editing by reducing background (Figs. 3b-e; Supplemental Figs. 6-8). The F22 variants of both inducible adenine base editors demonstrated a suitable dynamic range at the ABE16, ABE9, and HEK2 target sites, but high background in the absence of A115 was observed at the HEK3 locus (Figs. 3b-e). Higher background editing occurred only at the HEK3 target site for all editors tested, thus indicating that it is a locus-specific effect rather than a property of the nciCas9 switch.

**Figure 6.**
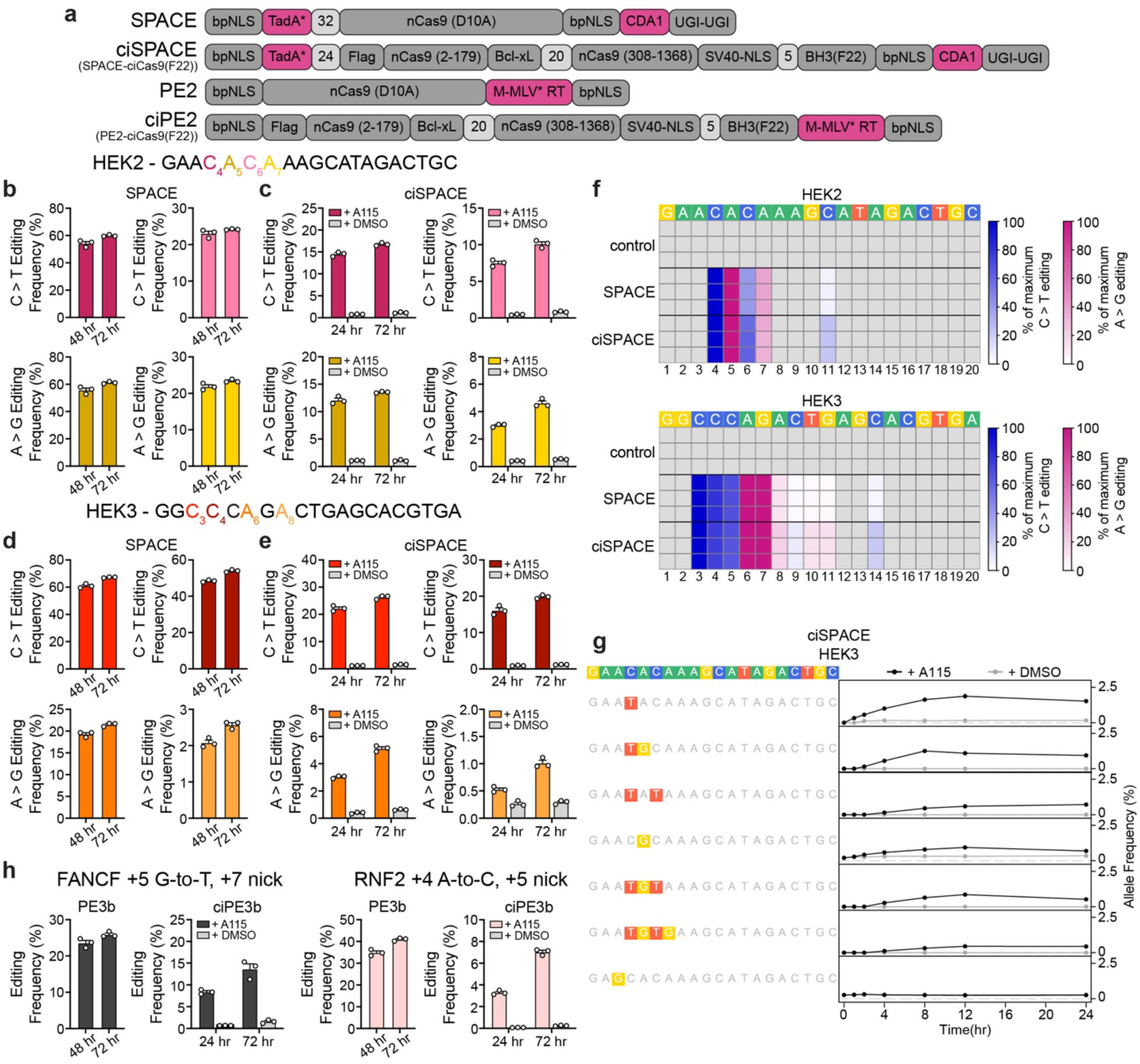
The ciCas9 switch can also be used to engineer chemically-controlled dual A-to-T and C-to-G base editors and prime editors. **a)** Domain arrangements of the unmodified SPACE base editor and PE2 prime editor and the chemically-controlled ciSPACE and ciPE2 editors. **b-e)** Dual A-to-T and C-to-G base editing by SPACE **(b,d)** and ciSPACE **(c,e)** at the HEK2 **(b,c)** and HEK3 **(d,e)** target sites. SPACE/ciSPACE base editing is shown at the 2 adenine and 2 cytidine nucleotides in each target site with the highest editing frequency with the Cas9 version of SPACE. The 4 different nucleotides in each target site are indicated by color in the target sequence. SPACE editing was quantified at 48 and 72 hr after cotransfection of base editor and sgRNA into HEK-293T cells. ciSPACE editing was quantified at 24 and 72 hr after 1 μM A115 or DMSO addition to HEK-293T cells. Bars show mean editing frequency ± SEM of 3 cell culture replicates with white circles showing individual replicates. **f)** Heatmaps of SPACE and ciSPACE editing through the entire HEK2 (top) and HEK3 (bottom) target sites. A-to-G base editing is shown in pink, C-to-T base editing is shown in blue. Editing is shown as a percentage of the highest edited nucleotide for each editor for that target site. Each row shows an individual cell culture replicate. SPACE editing frequencies were quantified at 72 hr after transfection and ciSPACE editing frequencies were quantified at 72 hr after 1 μM A115 addition to HEK-293T cells. The control shows untransfected cells harvested at the same time as ciSPACE. The numbers below the heatmaps show the position of the nucleotide from the most PAM-distal nucleotide. **g)** Time course of allele formation by ciSPACE at the HEK3 target sequence in HEK-293T cells treated with 1 μM A115 (black lines and circles) or DMSO. Cells were harvested and editing was quantified at specified time points after A115 or DMSO (gray lines and circles) addition Data represented as mean allele frequency ± SEM of 3 cell culture replicates. **h)** Prime editing frequencies by PE3b and ciPE3b at FANCF (black) and RNF2 (pink). PE3b editing frequencies were quantified at 48 and 72 hr after transfection and ciPE3b editing frequencies were quantified at 24 and 72 hr after 1 μM A115 addition to HEK-293T cells. Bars show mean editing frequency ± SEM of 3 cell culture replicates with white circles showing individual replicates.

We used the codon optimized chemically-controlled adenine base editors containing the F22 BH3 variant, ciABEmax and ciABE8e, for all subsequent experiments. ciABEmax and ciABE8e show similar editing windows as the parental versions (Figs. 3f-g; Supplemental Figs. 10c-d) with minimal editing activities in the absence of sgRNA (Supplemental Figs. 9e-h), minimal indel formation (Supplemental Figs. 11e-h), and low off-target editing activities (Supplemental Figs. 12c-d).

### Chemically-controlled base editors reveal how nucleotide position affects base editing kinetics

A key application of chemically-controlled enzymes, including Cas9, is exploring the kinetics of enzyme activity and downstream cellular processes using time course experiments. For example, chemically- and light-controlled CRISPR-Cas9 systems have been used to study the kinetics of DNA repair after DSB formation^26, 27, 29^. We previously found that the open ciCas9 switch was able to bind target sites and initiate DNA cleavage within minutes of A115 addition which can allow for precise interrogation of *in vivo* base editing kinetics^26^. Base editing time courses could provide insight into the relative kinetics of different DNA deaminase enzymes, reveal positional effects on editing kinetics at target sites that contain multiple editable nucleotides, and shed light on the relationship between deamination, repair, and editing. All four chemically-controlled base editors yielded appreciable editing within 24 hr of A115 addition, suggesting that activity is induced rapidly (Figs. 2b-g, 3b-e). Thus, for ciBE4max, ciABEmax and ciABE8e, we quantified editing at 1, 2, 4, 8, 12, and 24 hrs after A115 addition at 4 different target sites (Fig. 4; Supplemental Figs. 13-17)^40^. We did not evaluate ciAncBE4max due to its similarity to ciBE4max. For ciBE4max, the fastest-edited nucleotides within each target site began accumulating edits within 2-4 hr after activation (Fig. 4a; Supplemental Figs. 13a). Thus, within 2-4 hr of ciBE4max localization to a target site, a sufficient amount of deamination, DNA nicking, and DNA repair occurs to accumulate measurable base editing. Once a detectable level of editing was observed, base edits by ciBE4max accumulated nearly linearly for the first 12 hr at all four target sites (Supplemental Fig. 14a, 15). The rate of C-to-T base editing at different nucleotides within each target site correlated with the eventual magnitude of editing observed after 24 and 72 hr, with nucleotides in positions 5-7 (counting the PAM as positions 21–23) edited the earliest, fastest, and to the eventual greatest magnitude (Fig. 2h; Supplemental Figs. 10a, 14a, 15). Nucleotides outside the positions 5-7 demonstrated slower editing kinetics, which correlated with less overall editing at 24 and 72 hr. Thus, we found that the early editing kinetics of ciBE4max at each nucleotide position dictated the eventual editing magnitudes observed at later time points.

The chemically-controlled adenine base editors showed similar positional effects as ciBE4max in terms of early base editing kinetics and subsequent editing magnitudes (Figs. 3f-g, 4B-C; Supplemental Figs. 10c-d, 14b-c, 16, 17). For ciABEmax, the fastest edited nucleotide, usually at position 5, within each target site began accumulating base edits 1-2 hr after A115 addition, with early editing kinetics at all nucleotide positions correlating with eventual editing magnitudes at 24 and 72 hr (Figs. 3f, 4b; Supplemental Figs. 10c, 13b, 14b, 16). The HEK3 target site, where the adenine base editors had high background activity, showed accumulation of A115-promoted edits starting at later time points (Supplemental Figs. 13b). Like ciABEmax, ciABE8e also yielded A-to-G base edits at position 5 fastest, resulting in the greatest eventual magnitude at this position (Fig. 4c; Supplemental Figs. 13c). ciABE8e drove faster editing at 15 of 17 nucleotides across all target sites studied relative to ciABEmax, where the largest differences in kinetics were observed at positions that were poorly edited by ciABEmax (Supplemental Figs. 14b, c). Thus, the faster kinetics of ciABE8e resulted in editing across a broader window within a target site and lower selectivity for preferred positions (bystander editing) as compared to ciABEmax (Fig. 3g; Supplemental Fig. 10d). Thus, ciBE4max, ciABEmax, and ciABE8e enable *in vivo* kinetic studies, revealing that early editing kinetics correlate with the magnitude of editing later and highlighting the kinetic differences between deaminase enzymes which have previously only been explored *in vitro*^39, 41^.

To further investigate the positional effects of base editing kinetics, we normalized editing frequency at each position within every target site to the maximal editing at any position in the target site to allow comparisons across target sites (Fig. 4d-f). At nucleotides within the ciBE4max editing window, positional effects on the kinetics of base editing were readily apparent (Fig. 4d). Across all target sites, C-to-T editing by ciBE4max occured fastest at positions 5-7 (Fig. 4d), which was reflected in the greater magnitudes of editing achieved at positions 5-7 with ciBE4max at 72 hr (Fig. 2h; Supplemental Fig. 10a). ciABEmax and ciABE8e showed similar positional effects on editing rate and magnitude as ciBE4max, where adenines at positions 5 and 6 were edited the fastest (Figs. 4e, f). ciABE8e showed more rapid editing at positions 4, 7, and 8 compared to ciABEmax, emphasizing the broadened editing window of ciABE8e. Editing at positions 4 and 7 by ciABE8e was almost as fast as editing at positions 5 and 6. Furthermore, ciABE8e showed editing at positions 9 and 12 across multiple target sites, unlike ciABEmax. Thus, editing magnitude within a target site was dictated by the position of the substrate nucleotide: at every target site we tested, positions 5-7 were edited at high magnitudes due to rapid early editing kinetics whereas nucleotides both 3’ and 5’ of these rapidly edited positions were edited slower and thus to a lower magnitude.

### Chemically-controlled base editors provide insight into the kinetics of multiply-edited allele formation and nucleotide editing dependency

Analyzing the kinetics of cumulative editing at individual nucleotides within a target site provided insight into positional effects, but this approach masks the heterogeneity of base editing outcomes within individual cells. In particular, target sites with multiple A or C nucleotides are able to acquire different combinations of multiple edits, resulting in the accumulation of different alleles. The longer the period of active base editing, the greater the accumulation of these multiply-edited alleles^42^. Using our chemically-controlled base editors, we, for the first time, dissected the kinetics of multiply-edited allele formation *in vivo* to better understand the order in which nucleotides are edited and the impact of initial edits on subsequent ones in the formation of multiply-edited alleles.

To determine the order in which nucleotides are edited within a target site, we identified all distinct combinations of edits (i.e. alleles) at four target sites for the ciBE4max, ciABEmax, and ciABE8e base editors and tracked the frequency of each allele over time (Fig. 5a-c, Supplemental Figs. 18-20). As expected, we observed early accumulation of singly-edited alleles and later accumulation of multiply-edited alleles. Singly-edited alleles began to accumulate within 1-2 hr, similar to the time frame observed in the cumulative nucleotide editing analysis (Figs. 5a-c; Supplemental Fig. 14). Generally, all alleles accumulated linearly for 2-6 hr after they were first detected. However, some alleles eventually decreased in accumulation rate or even in frequency. We hypothesized that these decreases reflected consumption of these alleles to form more highly edited alleles. For example, the singly-edited A5G allele created by ciABEmax at ABE16 decreased in accumulation rate between 4 and 8 hr and then decreased in frequency after 8 hr (Fig. 5b). Presumably, this A5G allele was being consumed to form the doubly-edited alleles A5G/A7G or A4G/A5G, which both began to accumulate 4 hr after activation. The triply-edited A4G/A5G/A7G allele then appeared later, first at 8 hr following a likely third edit of the A4G/A5G or A5G/A7G alleles. We found that, in all cases, alleles with fewer edits appeared first, followed by alleles with more edits.

The faster editing kinetics and larger range of positions edited by ciABE8e compared to ciABEmax is also reflected in the greater diversity and frequency of higher order edited alleles detected at all four target sites (Figs. 5b-c; Supplemental Figs. 19-20). For example, the A4G/A5G/A7G allele at ABE16 accumulated linearly with ciABEmax to a maximum frequency of 3.2% at 24 hr (Fig. 5b). With ciABE8e, the A4G/A5G/A7G allele appeared at a frequency of 4.5% within 8 hr whereupon accumulation slowed, presumably due to the consumption of this allele to form higher order alleles (Fig. 5c). Furthermore, for ciABE8e, we rarely detected singly-edited alleles, suggesting that singly-edited alleles were quickly consumed to form higher order alleles. Thus, the faster *in vivo* editing kinetics of ciABE8e compared to ciABEmax results in the generation of higher order alleles rather than a greater frequency of lower order alleles. Multiply-edited alleles can be explained by two kinetic models. An independent model posits that editing at one position does not impact the rate of editing at other positions within a target site in the formation of a multiply-edited allele. A dependent model posits that editing at one position affects the rate of editing at other positions. Under the independent model, the frequency of a multiply-edited allele at a particular time should be the product of the frequency of the individual edits it contains at that time. For each base editor at each target site, we computed the observed single edit frequencies from our cumulative editing analysis (Supplemental Fig. 14). Then, we computed the expected frequency of each multiply-edited allele by multiplying the frequencies of the constituent single edits. We compared the expected and observed frequencies for each allele for all time points in which that allele was detected. If the expected allele frequency is equal to or greater than the measured frequency over all time points where the allele is detected, we classified that allele as “independent.” If the expected allele frequency was less than the measured frequency over all time points where the allele is detected, we classified that allele as “dependent.” Across all base editors and target sites, 28 of 31 multiply-edited alleles had an expected frequency that was less than the measured frequency at all time points where the allele was detected (Fig. 5d; Supplemental Figs. 21-23). 27 of 28 dependent alleles showed statistical significance in dependence with a permutation analysis based on the Chi-squared test statistic (Materials and Methods; Supplemental Table 3). Thus, these alleles were dependent, suggesting that editing of the first nucleotide increased the rate of editing at all subsequent nucleotides. The remaining three alleles initially appear to be dependent, with measured frequencies higher than expected, but show decreased allele accumulation at later time points compared to the expected allele accumulation (Fig. 5e). Therefore, these alleles cannot be classified as either independent or dependent using our expected allele frequency analysis, and we thus classify them as ambiguous.

### The ciCas9 switch can also be used to engineer chemically-controlled dual A-to-T and C-to-G base editors and prime editors

Given that the ciCas9 switch provides chemical control of transcriptional activation and cytidine and adenine base editors by modulating DNA binding, we wondered whether the switch could also be applied to dual A-to-T and C-to-G base editors and to prime editors^8, 13^. One of the reported dual A-to-T and C-to-G base editors, SPACE, utilizes two deaminase domains fused to nCas9^8^. When SPACE binds to a target site through the nCas9 domain, it can create both A-to-T and C-to-G base edits within a single target site. We generated a chemically-controlled dual A-to-T and C-to-G base editor using the nciCas9 switch, ciSPACE, constructed with the same domain architecture as the unmodified version (Fig. 6a). We found that ciSPACE was able to introduce both C-to-T and A-to-G edits at an sgRNA-defined target site upon A115 addition in cells transiently expressing ciSPACE, with minimal background editing (Fig. 6b-e). Moreover, ciSPACE edits at the exact same positions within target sites as SPACE (Fig. 6f). ciSPACE also forms minimal indels and off-target base edits, at magnitudes similar to or lower than SPACE (Supplemental Figs. 24-25). We next explored the kinetics of the two deaminase domains with time course experiments (Fig. 6g, Supplemental Fig. 26). At all three target sites investigated, C-to-T edits appeared to accumulate faster than A-to-G edits. Thus, at least at these target sites, cytidine deamination and repair appear much faster than adenine deamination and repair.

Finally, we applied the nciCas9 switch to the PE2 prime editor enzyme, which consists of the nCas9(H840A) variant fused to M-MLV reverse transcriptase used in combination with pegRNA/sgRNA pairs to effect base substitutions and small insertions or deletions^13^. We constructed a chemically-controlled PE2 enzyme, ciPE2, in the same domain architecture as the unmodified version (Fig. 6a). We tested two sets of previously reported pegRNA/sgRNA pairs with ciPE2 and observed incorporation of the desired edit, albeit less efficiently than the PE2 editor (Fig. 6h). Moreover, we observe minimal indel formation at the prime editing target site, similar to that of the PE2 editor (Supplemental Fig. 27). Thus, the ciCas9 switch can be applied to Cas9-based effectors with diverse architectures by simply replacing Cas9 with ciCas9, including to control dual base editing and prime editing with minimal unwanted editing.

## Discussion

We demonstrate a general method for gaining precise chemical control over different Cas9-based effectors by modulating DNA target site binding using the ciCas9 switch. Because the ciCas9 switch consists only of the replacement of the Cas9 REC2 domain with Bcl-xL and appendage of a BH3 peptide, it can be installed while preserving nearly any desired Cas9-based effector architecture. Use of the ciCas9 switch to engineer chemically-controlled transcriptional activators and base editors was simple compared to currently available multi-protein systems that require screening for functional split proteins and careful co-expression of multiple protein components^24, 30–34^. Rapid activation kinetics mean the ciCas9 switch is also more temporally precise than other chemically-controlled Cas9 systems that rely on relocalization of protein to the nucleus or shutoff of degradation, processes that have much slower kinetics^17–23^. As a result of using domain replacement to confer chemical control over Cas9 activity, the overall size of the ciCas9 switch is similar to that of Cas9 itself. Furthermore, many different Bcl-xL/BH3 disruptors can be used to activate the ciCas9 switch and are compatible with a variety of organisms^43, 44^. Thus, ciCas9 can easily replace Cas9 in any Cas9-based effector to confer chemical control over effector activities. We used the ciCas9 switch to gain chemical control of transcriptional activation, base editing, and prime editing, demonstrating the versatility and simplicity of the switch.

The high temporal precision of the ciCas9 switch allowed us to obtain unique insight into *in vivo* base editing kinetics. Using three chemically controlled base editors, ciBE4max, ciABEmax and ciABE8e, we elucidated the early *in vivo* kinetics of base editing. Rapid early editing 5-7 nucleotides from the 5’ end of the target site generally led to higher editing at later time points. Investigation of the kinetics of multiply-edited alleles revealed that they do not form following an independent model of editing. Instead, bystander edits form at an increased rate following a single editing event at a preferred position. We hypothesize that editing dependency arises from one of two different mechanisms depending on whether the base editor remains bound to the target site during the entire editing process or undergoes cycles of dissociation and rebinding. If the base editor remains bound, then the first base edit within a target site likely increases the accessibility of bystander nucleotides within the same target site to the deaminase yielding faster bystander editing. If the base editor undergoes cycles of dissociation and rebinding, the first deamination event would create a mismatch in the DNA double helix, thus favoring subsequent cycles of base editor rebinding and deamination. Discriminating between these possibilities requires investigation of the *in vivo* kinetics of binding and dissociation as well as direct measurement of deamination rather than editing to dissect deaminase activity and subsequent DNA repair. Insights into the kinetics of base editing, and, especially, multiply-edited allele formation could inform future efforts to engineer more efficient and selective base editors, which are sorely needed for precise correction of pathogenic mutations at target sites containing bases that could be unintentionally edited^45^ and for pooled screening of DNA variant effects of genes at their endogenous locus^46^ where the unpredictable and partially specific nature of base editing complicates assessment of DNA variants without sequencing the edited locus.

Despite the utility and power of the ciCas9 switch, challenges remain. Installing the ciCas9 switch often modestly decreases the efficiency of Cas9-based effector. We also observe appreciable DMSO background for the chemically-controlled base editors at select target sites, which we mitigated by increasing the strength of the Bcl-xL/BH3 peptide autoinhibitory switch^47^. While some background editing remains and could be problematic for therapeutic applications, low background editing is observed prior to ciCas9 switch activation in our time course experiments and provides ample dynamic range to allow insight into editing kinetics. Additionally, our experiments did not fully capture the uridine and inosine base editing intermediates. We used Kapa HiFi polymerase, which is inefficient at amplifying DNA templates containing a uridine, and inosine bases can be read as any DNA base with cytidine-inosine base pairing being the most efficient^48, 49^. Thus, the base editing activities we report may be an underestimate. Furthermore, development of assays to directly measure deaminase activity and DNA repair *in vivo* coupled with computational modeling of the data is needed to provide a more accurate picture of base editing mechanisms and the timing of different allele outcomes. With these tools in hand, ciCas9 base editors could be used to generate data to help improve the recently reported model of bystander base editing^50^. These tools, along with ciCas9 base editors could also be used to develop editors with desirable kinetic properties and reduced bystander editing activity. However, we revealed that merely changing the overall rate of editing is not enough to develop more selective base editors, because bystander editing is a result of a dependent process.

We have shown that the ciCas9 switch offers a general approach to engineering chemical control of Cas9-based effectors. For example, the ciCas9 switch could be used to temporally control the expression of specific genes during different stages of development or cell differentiation. Precise definition of a starting time for lineage tracing is also achievable with the ciCas9 switch. Temporal control over base or prime editing of clinically relevant loci could also be beneficial to better control editing efficiency and specificity. Finally, other Cas9-based effector proteins could be temporally controlled using the ciCas9 switch such as directly reading or writing chromatin marks and colocalizing genetic elements within the genome to study the effects of 3D genome architecture. We envision that the ciCas9 switch can be applied to confer temporal control over a plethora of Cas9-based effector proteins that are currently available or will be engineered in the future.

## Methods

### Expression plasmids

Mammalian expression plasmids of dCas9, dciCas9(L22), and dciCas9(L22)-VPR were expressed using the pcDNA5/FRT/TO backbone (ThermoFisher). dciCas9(L22) was constructed by introducing the D10A and H935A (dCas9 H840A equivalent) mutations into previously reported ciCas9(L22)^26^. To create dciCas9(L22)-VPR, PCR-amplified VP64-p65-Rta from pEF045, a gift from Jesse Zalatan, was assembled with dciCas9(L22) into the pcDNA5/FRT/TO backbone linearized using the BamHI and EcoRV restriction digest sites. dCas9-VPR was expressed in a pEF backbone and was a gift from David Shechner. The expression of MCP-VPR and CXCR4 scRNA were from a single vector containing both U6 and CMV promoters, a gift from Jesse Zalatan.

All ciCas9 base editors and prime editors were expressed using the pcDNA3.1(+) backbone. nciCas9(D10A) and nciCas9(H840A) were constructed using PCR amplification products from previously constructed ciCas9(L22) and dciCas9(L22). Codon optimized ciCas9(L22) and dciCas9(L22) were purchased from Twist Biosciences. The Bcl-xL and BH3 components were codon optimized using the Genscript codon optimization tool. nciCas9(F22) constructs were made by introducing a point mutation into the L22 constructs using Gibson assembly. The deaminase and UGI components in ciBE4max and ciAncBE4max were PCR-amplified from pCMV_BE4max (Addgene #112093) and pCMV_AncBE4max (Addgene #112094), respectively, both gifts from David Liu. The deaminase components in ciABEmax and ciABE8e were PCR-amplified from pCMV_ABEmax (Addgene #112095) and pCMV_ABE8e (Addgene #138489), respectively, both gifts from David Liu. The M-MLV* reverse transcriptase in ciPE2 was PCR-amplified from pCMV_PE2 (Addgene #132775), a gift from David Liu. The base and prime editing components were assembled with the nciCas9(L22/F22) component using Gibson assembly into the pcDNA3.1(+) backbone linearized using the BamHI and EcoRI restriction digest sites. For a full list of constructs and corresponding amino acid sequences, see Supplemental Table 4.

All sgRNAs were cloned into the gRNA cloning vector (Addgene #41824), a gift from George Church. The CXCR4 sgRNA plasmid has been previously reported^31^. Briefly, a single-stranded DNA (ssDNA) oligo with overlap to the gRNA cloning vector 5’ and 3’ of the 20 nt target sequence was purchased from Integrated DNA Technologies (IDT). The ssDNA oligo was assembled into the gRNA cloning vector linearized using the AflII site by Gibson assembly.

### Mammalian cell culture

HEK-293T cells (ATCC, CRL-3216) and HEK-293 TREx FlpIn cells (ThermoFisher) were cultured in high glucose Dulbecco’s Modified Eagle Medium (DMEM, Gibco) supplemented with 10% (v/v) fetal bovine serum (FBS, MilliporeSigma). Cells were all incubated at 37 °C, 5% CO_2_ and found to be free from mycoplasma at least every 6 months. HEK-293 TREx FlpIn EMX1-EGFP reporter cells were cultured in DMEM supplemented with 10% (v/v) FBS and 50 μg/mL hygromycin (Mirus).

### EGFP reporter construction for dciCas9-VPR transcriptional activation

A 20 bp EMX1 target and PAM sequence and 20 bp of endogenous gDNA sequence 5’ and 3’ of the target were cloned using Gibson assembly into a pcDNA5-FRT-TO backbone lacking the CMV promoter. 3’ of the target sequence is a minimal promoter followed by 100 bp of random DNA sequence and an EGFP reporter gene (Supplemental Fig. 3)^51^. The plasmid containing this locus was then transfected into HEK-293 TRex Flp-In cells (ThermoFisher) along with a pOG44 plasmid encoding a Flp-recombinase using Turbofectin according to manufacturer’s protocols. Cells with successful integration of the reporter locus were selected using hygromycin (Mirus Bio).

### Transcriptional activation

For CXCR4 transcriptional activation with dciCas9-VPR, 6 × 10^4^ HEK-293T cells were seeded in 12-well plates. ∼20-24 hr after seeding cells, each well was transfected with 1 μg total dciCas9-VPR and CXCR4 sgRNA plasmids (450 ng dciCas9-VPR, 450 ng CXCR4 sgRNA, 100 ng mCherry control) using Turbofectin (Origene) according to the manufacturer’s protocol. ∼24 hr after transfection, a final concentration of 1 μM of A-1155463 (A115; ChemieTek) was added to the cells, final [DMSO] of 0.1%. 48 hr after A115 addition, cells were harvested and incubated with APC anti-human CD184 (CXCR4) [12G5] (BioLegend) for 1 hr and then fluorescence was analyzed on the LSRII flow cytometer (BD Biosciences). 30,000 single cell events were collected for each sample. The median APC fluorescence is reported for the brightest 25% of cells expressing mCherry transfection control. A similar protocol was followed for dciCas9 + scRNA transcriptional activation of CXCR4 with the exception of 450 ng dciCas9, 450 ng CXCR4 scRNA/MCP-VPR-IRES-mCherry, and 100 ng empty pcDNA5/FRT/TO plasmid was transfected into HEK-293T cells. All flow cytometry data was analyzed using FlowJo. See Supplemental Fig. 28 for example cell gating strategies.

For EGFP reporter transcriptional activation, a similar protocol as CXCR4 activation was used except an EMX1 sgRNA was used to target dciCas9-VPR and no antibody incubation was performed, cells were directly analyzed for EGFP fluorescence by flow cytometry. A115 was diluted to the indicated concentrations using DMSO and added to cells with a final [DMSO] of 0.1%. See Supplemental Fig. 29 for example cell gating strategies.

### Base editing and prime editing with Cas9 and ciCas9 base editors

For both base editing and prime editing experiments, HEK-293T cells were seeded at 1.8-2.0 × 10^4^ cells per well in a 12-well plate. ∼20-24 hr after seeding cells, cells were transfected with 1 μg total plasmid DNA of base/prime editor, sgRNA, and a pMAX-GFP transfection control using Turbofectin according to the manufacturer’s protocol. For base editing, 690 ng base editor, 230 ng sgRNA, and 80 ng of pMAX-GFP were cotransfected into each well. No sgRNA control experiments were conducted with 690 ng base editor and 310 ng of pMAX-GFP. For prime editing, 630 ng prime editor, 210 ng pegRNA, 70 ng sgRNA, and 90 ng of pMAX-GFP were cotransfected into each well. ∼24 hr after transfection, a final concentration of 1 μM of A115 was added to the wells containing ciCas9 base or prime editor, final [DMSO] of 0.1%. Cas9 base and prime editor conditions were harvested at the indicated time points after transfection. ciCas9 base and prime editor conditions were harvested at the indicated time points after A115 addition.

### Library preparation for targeted amplicon DNA sequencing

Genomic DNA isolation, sequencing, and indel frequency analysis for non-library loci were performed as previously described^35^. Briefly, genomic DNA was extracted from cells using the DNeasy Blood and Tissue kit (Qiagen) according to the manufacturer’s protocol with an extended proteinase K digestion of 1 hr at 56 °C. The loci of interest were first amplified with 15 cycles of PCR from 2 μL (∼100 ng) of genomic DNA eluate using a 5 μL Kapa HiFi HotStart polymerase reaction (Roche). The first PCR was then diluted with 25 μL of DNAse-free water. Indexes and Illumina cluster generation sequences were then added with a secondary PCR reaction using 3 μL of the diluted primary PCR product with a 10 μL Kapa Robust HotStart polymerase reaction (Roche) for 20 cycles. The final amplicons were run on a 1% TBE-agarose gel and DNA was extracted using a Freeze and Squeeze column according to the manufacturer’s protocol (BioRad). Gel extracted amplicons were quantified using the Qubit dsDNA HS assay kit (Invitrogen). Up to 1000 indexed amplicons were pooled and sequenced on a NextSeq 550 using a NextSeq Mid 150 v2/v2.5 kit (Illumina). A minimum of 1,000 reads was acquired for each sample except for replicate 1 of ciABE8e at ABE16 with 24 hr DMSO treatment in the time course experiments (Fig. 5c, Supplemental Fig. 16).

### Editing quantification and analysis

Editing was quantified using the CRISPResso2 package, version 2.0.45^52^. All editing was quantified using batch analysis. Base editing was quantified using the additional flags “-wc -10 - w 10 -q 30”. Cumulative base edits at each nucleotide within the target site were extracted from the output table “Nucleotide_percentage_summary.” Normalized base editing in Fig. 4d-f was calculated by setting the mean of the highest edited nucleotide within each target site at 24 hr to 100% editing. Normalized editing frequencies at other nucleotide positions and at other time points within the same target site were calculated as a percentage of the maximum editing at that highest edited nucleotide using the mean editing frequency of a triplicate of cell culture replicates. Allele frequencies were extracted from the output table “Alleles_frequency_table_around_sgRNA.” Allele frequencies were also determined by only looking at base changes within the 20 nucleotide target sequence, any changes outside the target sequence were trimmed and allele frequencies were summed using the custom script “allele_frequency_merge_v1.py.” Plotting of allele frequency time courses (Figs. 5a-c, 6g; Supplemental Figs. 18-23, 26) were filtered for alleles that were detected at >0.3% at any time point and for alleles that showed only A-to-G or C-to-T base edits corresponding to the base editor studied. 0.3% was the lowest threshold to filter out alleles that contained sequencing errors and non-A-to-G or non-C-to-T base edits. 0% editing frequency was imputed for alleles that were not detected at certain time points but showed >0.3% editing frequency at other time points.

Indel frequencies from the base editors were calculated using the output table “CRISPRessoBatch_quantification_of_editing_frequency” by calculating:

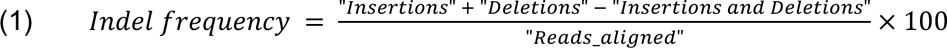

from the table columns. Heatmaps showing base editing frequencies were filtered to only show base conversion frequencies at A or C nucleotides within the target site. Editing frequencies in Figs. 2h-i, 3f-g, 6f, and Supplemental Fig. 10 were filtered for positions with base conversion greater than 0.7%. Heatmaps showing off-target base editing in Supplemental Fig. 12 were filtered for positions with base conversion greater than 0.1%.

Prime editing was quantified using the additional flag “-q 30”. Prime editing frequencies were calculated using the output table “CRISPRessoBatch_quantification_of_editing_frequency” using the “Prime-edited” row for each sample and calculating:

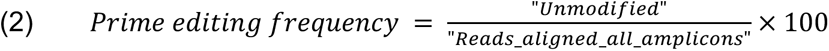

from the table columns. Indel frequencies from prime editing were analyzed using standard NHEJ CRISPResso2 settings. Indel frequencies were then calculated using the output table “CRISPRessoBatch_quantification_of_editing_frequency” using the equation (1) and the same table columns.

For analysis of editing at early time points in Figs. 4a-c and Supplemental Fig. 13, statistical comparison of editing at 0 hr to 1, 2, and 4 hr after A115 addition to editing was completed using a One-way ANOVA using the Graphpad Prism 9 software. Results from the One-way ANOVA are reported in Supplemental Table 2.

### Calculation of expected allele frequencies

The expected frequency for an allele was calculated as a product of the frequency of edits at each nucleotide position that make up the allele:

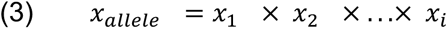

Where *x*_i_ represents the edited nucleotide frequency at nucleotide position *i*. The relative error was calculated using the standard error of the mean for each nucleotide position:

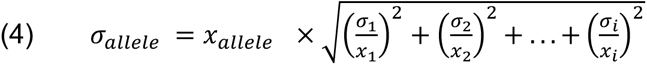

Where *σ*_i_represents the standard error of the mean at nucleotide position *i*.

To determine the dependent vs. independent models of editing for each allele, we compared expected versus measured allele frequencies at all time points where the allele was detected.

A permutation analysis based on the Chi-squared test statistic was used to identify alleles with a measured frequency that is significantly higher than their expected frequency. The cumulative frequency for each nucleotide within an allele for this analysis was calculated by summing the frequency at which an edit at a specific nucleotide appears as a singleton or in combination with other edited nucleotides. The chi-squared statistic was normalized by the number of time points in which the expected frequency was >0. We classified alleles with a chi-squared statistic >0.045 in at least two of three replicates as dependent. This threshold was determined using a background distribution generated by shuffling the data (since not all the chi-squared test assumptions hold in this case).

## Acknowledgements

This work was supported by the NIH, grant no. RM1HG010461 (NGHRI) and R01GM109110 (NIGMS) to D.M.F., and grant no. R01GM145011 (NIGMS) to D.J.M. O.P. was supported in part by a fellowship from the Edmond J. Safra Center for Bioinformatics at Tel-Aviv University. E.B. is a Faculty Fellow of the Edmond J. Safra Center for Bioinformatics at Tel Aviv University.

## Author contributions

C.T.W., D.J.M., and D.M.F. conceived of the work and wrote the manuscript. C.T.W. performed all experiments. O.P. and E.B. performed the permutation analysis based on the Chi-squared test statistic for the base editing dependency analysis.

## Data and code availability

Sequencing data is available in the Sequence Read Archive (SRA) under accession number #########. Scripts written for parsing data and plotting figures are available on Github (https://github.com/cindytxwei/ciCas9effectors). Scripts written for the permutation analysis based on the Chi-squared test statistic are also available on Github (https://github.com/omripel/BEAnalysis).

## SUPPLEMENTAL FIGURES

**Supplemental Figure 1.**
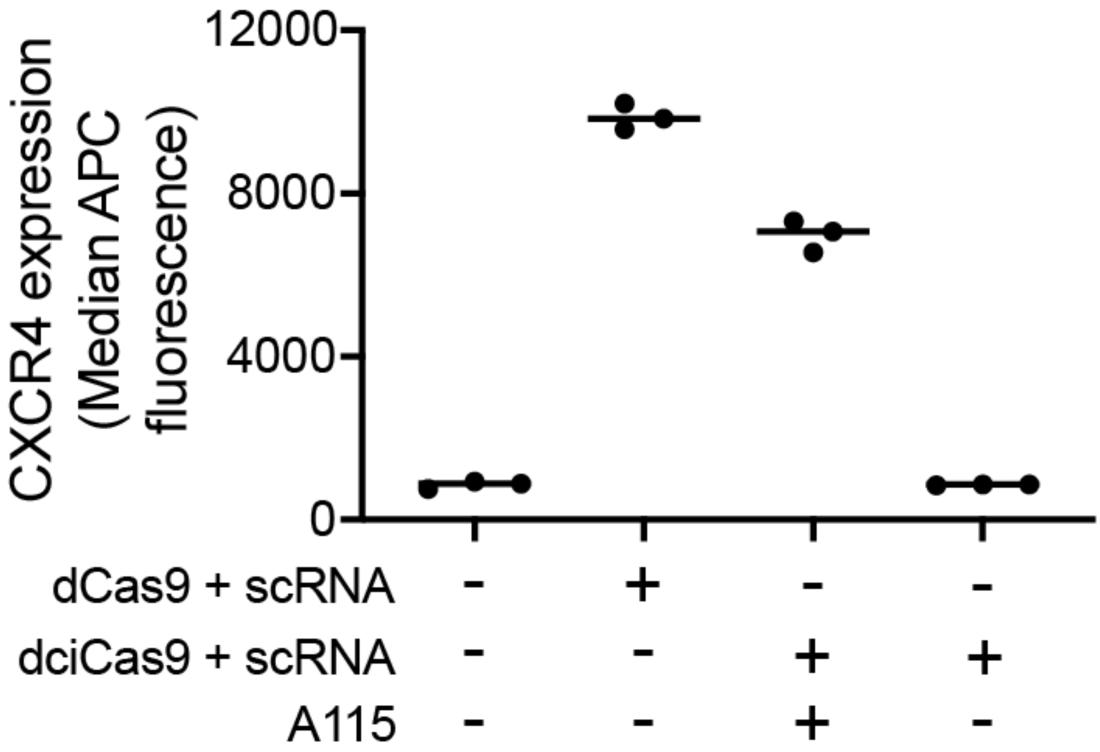
dciCas9 transcriptional activation using a scRNA. Activation of CXCR4 expression using dCas9 or dciCas9 with a scRNA targeted to the promoter region in HEK-293T cells that recruits MCP-VPR in the presence or absence of 1 μM A115 activation for 48 hr. Data represented as 3 cell culture replicates shown with a line showing the mean.

**Supplemental Figure 2.**
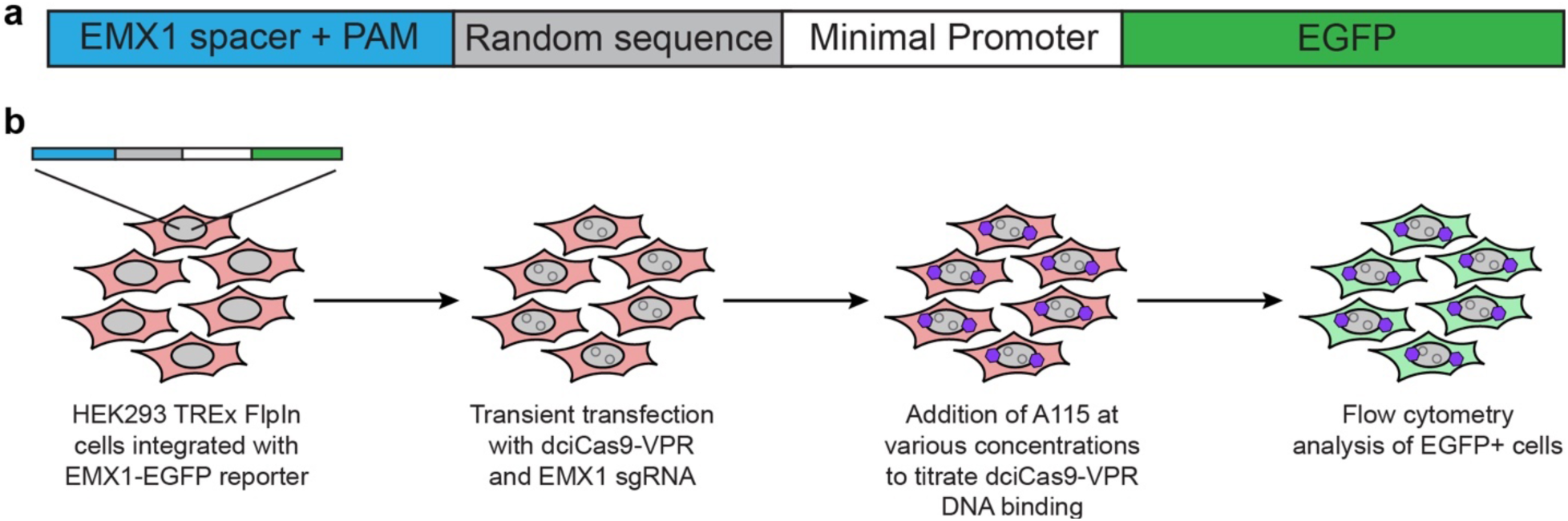
Schematic of EMX1-EGFP reporter locus in HEK-293 TREx FlpIn cells. **a)** Schematic of the EMX1-EGFP transcriptional synthetic locus integrated into HEK-293 TREx FlpIn cells. **b)** Workflow of using EMX1-EGFP transcriptional synthetic locus cells with dciCas9-VPR + EMX1 sgRNA for a dose-response of EGFP expression.

**Supplemental Figure 3.**
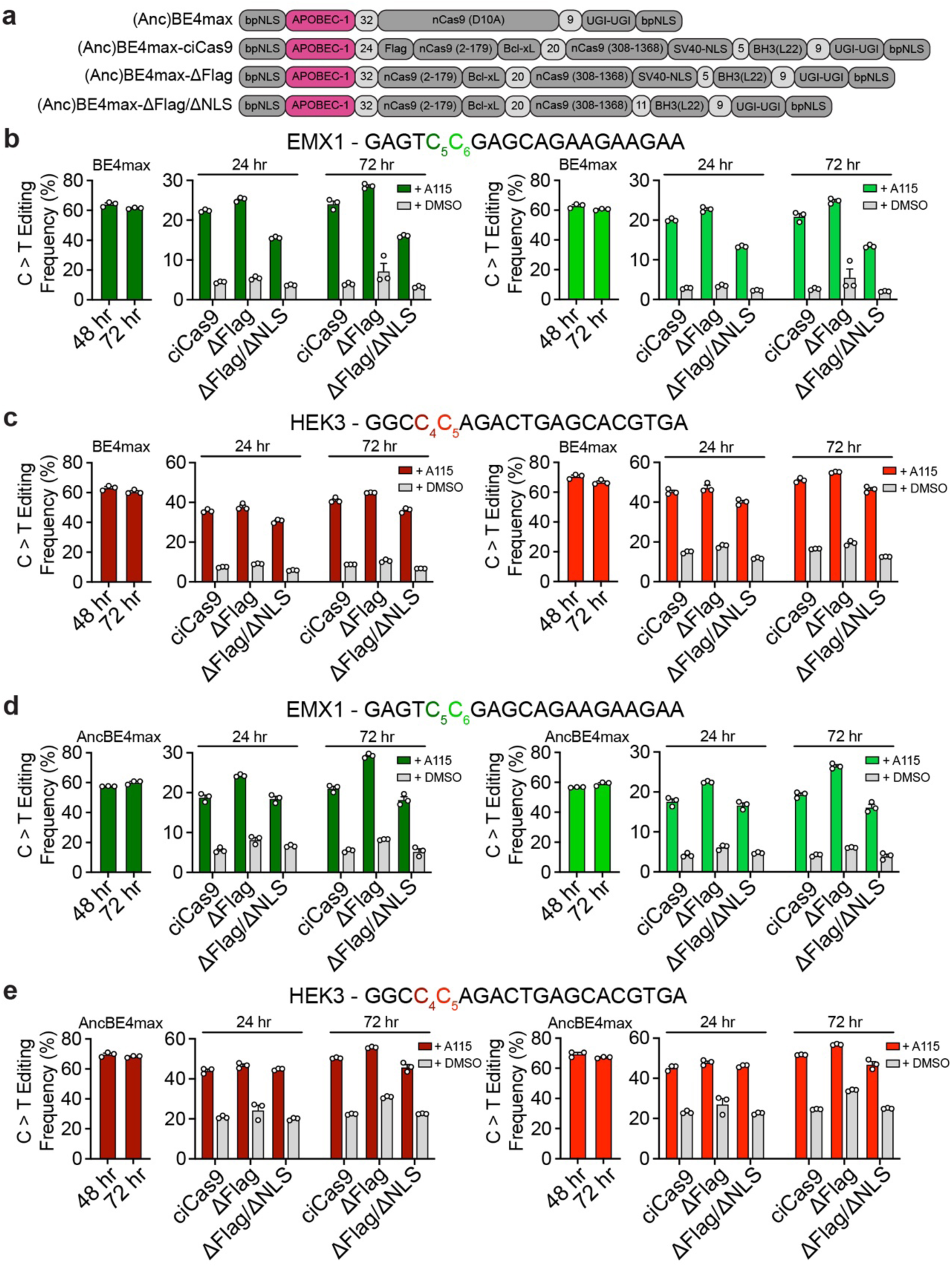
Chemically-controlled cytidine base editors without codon optimization. **a)** Schematic of the domain arrangements in the unmodified BE4max and AncBE4max base editors and the chemically-controlled BE4max and AncBE4max base editors without codon optimization and using the ciCas9(L22) variant. 3 different versions of ciCas9 were used, ciCas9(L22), ciCas9(L22) without a Flag-tag (ΔFlag), and ciCas9(L22) without a Flag-tag and additional SV40-NLS (ΔFlag/ΔNLS). **b-c)** C-to-T editing frequency with BE4max and BE4max-ciCas9 at the EMX1 **(b)** and HEK3 **(c)** target sites. **d-e)** C-to-T editing frequency with AncBE4max and AncBE4max-ciCas9 at the EMX1 **(d)** and HEK3 **(e)** target sites. BE4max and AncBE4max editing were measured at 48 and 72 hr after co-transfection of BE4max and sgRNA. BE4max-ciCas9 and AncBE4max-ciCas9 editing were measured at 24 and 72 hr after 1 μM A115 addition. C-to-T editing is shown at the 2 nucleotides in each target site with highest editing frequency with the Cas9 version of base editors (BE4max or AncBE4max). The 2 different nucleotides are indicated by color in the target sequence. Bars show mean editing frequency ± SEM of 3 cell culture replicates with white circles showing individual replicates.

**Supplemental Figure 4.**
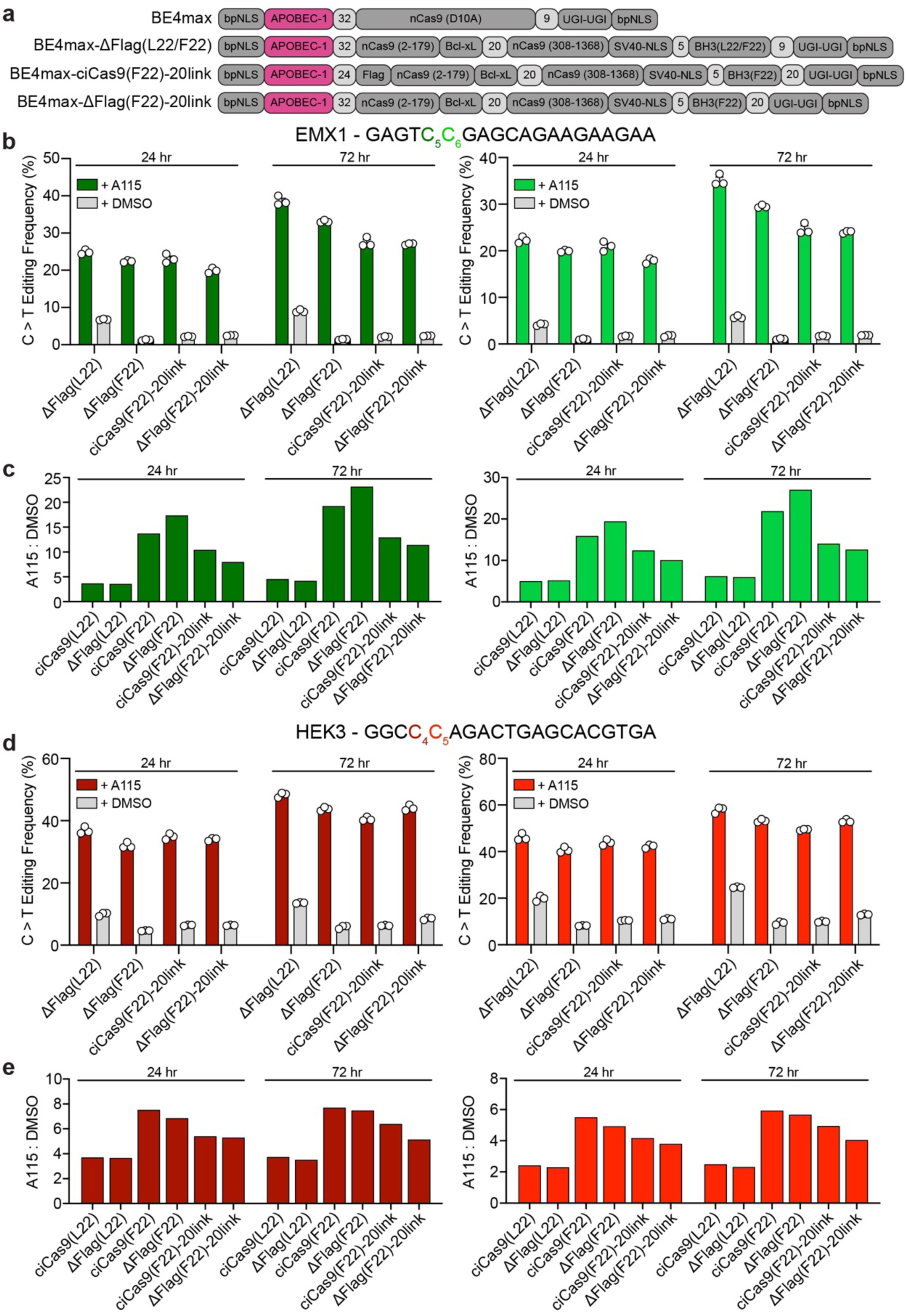
Additional constructs of codon optimized chemically-controlled BE4max editors. **a)** Schematic of domains in the unmodified BE4max base editor and additional constructs of the codon optimized BE4max-ciCas9 base editors tested. 4 different versions of ciCas9 were additionally tested: No-Flag-ciCas9(L22) (ΔFlag(L22)), No-Flag-ciCas9(F22) (ΔFlag(F22)), ciCas9(F22)-20linker-2xUGI (ciCas9(F22)-20link), and No-Flag-ciCas9(F22)-20linker-2xUGI (ΔFlag(F22)-20link). **b,d)** C-to-T editing frequencies of the BE4max-ciCas9 constructs at the EMX1 **(b)** and HEK3 **(d)** target sites. C-to-T editing is shown at the 2 nucleotides in each target site with highest editing frequency with BE4max. The 2 different nucleotides are indicated by color in the target sequence. Editing by all BE4max-ciCas9 constructs are quantified at 24 and 72 hr after 1 μM A115 or DMSO addition to HEK-293T cells. Bars show mean editing frequency ± SEM of 3 cell culture replicates with white circles showing individual replicates. **c,e)** Ratio of the mean C-to-T editing frequency with 1 μM A115 to the mean C-to-T editing frequency with DMSO (A115:DMSO) for all tested BE4max-ciCas9 base editors in Fig. 2 and Supplemental Fig. 4 at the EMX1 **(c)** and HEK3 **(e)** target sites. Bars show the ratios of editing at the 2 nucleotides at each target site with highest editing frequency with BE4max. Editing frequencies used to calculate the ratio were measured at 24 and 72 hr after A115 addition to HEK-293T cells.

**Supplemental Figure 5.**
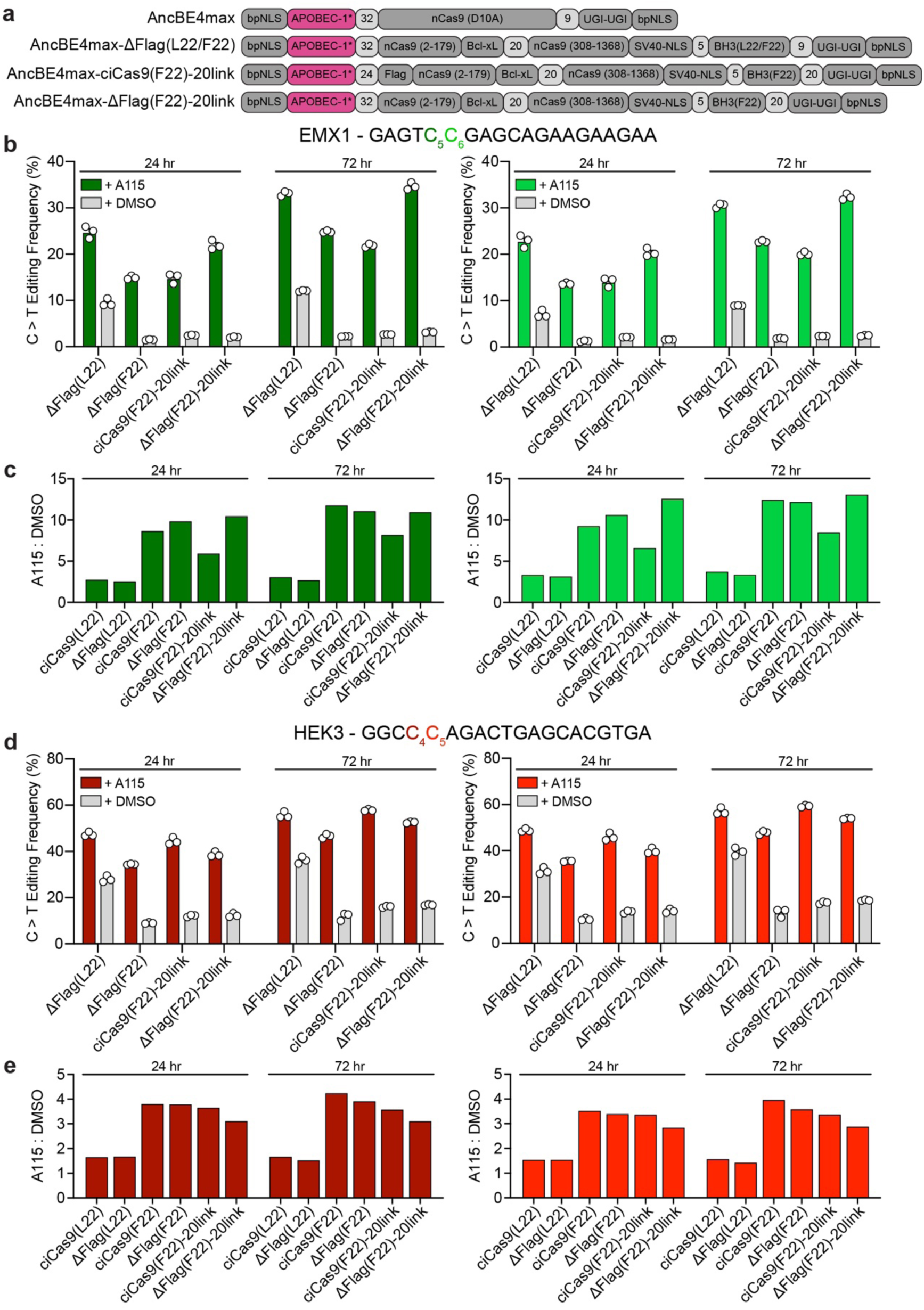
Additional constructs of codon optimized chemically-controlled AncBE4max editors. **a)** Schematic of domains in the unmodified AncBE4max base editor and additional constructs of the codon optimized AncBE4max-ciCas9 base editors tested. 4 different versions of ciCas9 were **b,d)** C-to-T editing frequencies of the AncBE4max ciCas9 constructs at the EMX1 **(b)** and HEK3 **(d)** target sites. C-to-T editing is shown at the 2 nucleotides at each target site with highest editing frequency with AncBE4max. The 2 different nucleotides are indicated by color in the target sequence. Editing by all AncBE4max-ciCas9 constructs are quantified at 24 and 72 hr after 1 μM A115 or DMSO addition to HEK-293T cells. Bars show mean editing frequency ± SEM of 3 cell culture replicates with white circles showing individual replicates. **c,e)** Ratio of the mean C-to-T editing frequency with 1 μM A115 to the mean C-to-T editing frequency with DMSO (A115:DMSO) for all tested AncBE4max-ciCas9 base editors in Fig. 2 and Supplemental Fig. 5 at the EMX1 **(c)** and HEK3 **(e)** target sites. Bars show the ratios of editing at the 2 nucleotides in each target site with highest editing frequency with AncBE4max. Editing frequencies used to calculate the ratio were measured at 24 and 72 hr after A115 addition to HEK-293T cells.

**Supplemental Figure 6.**
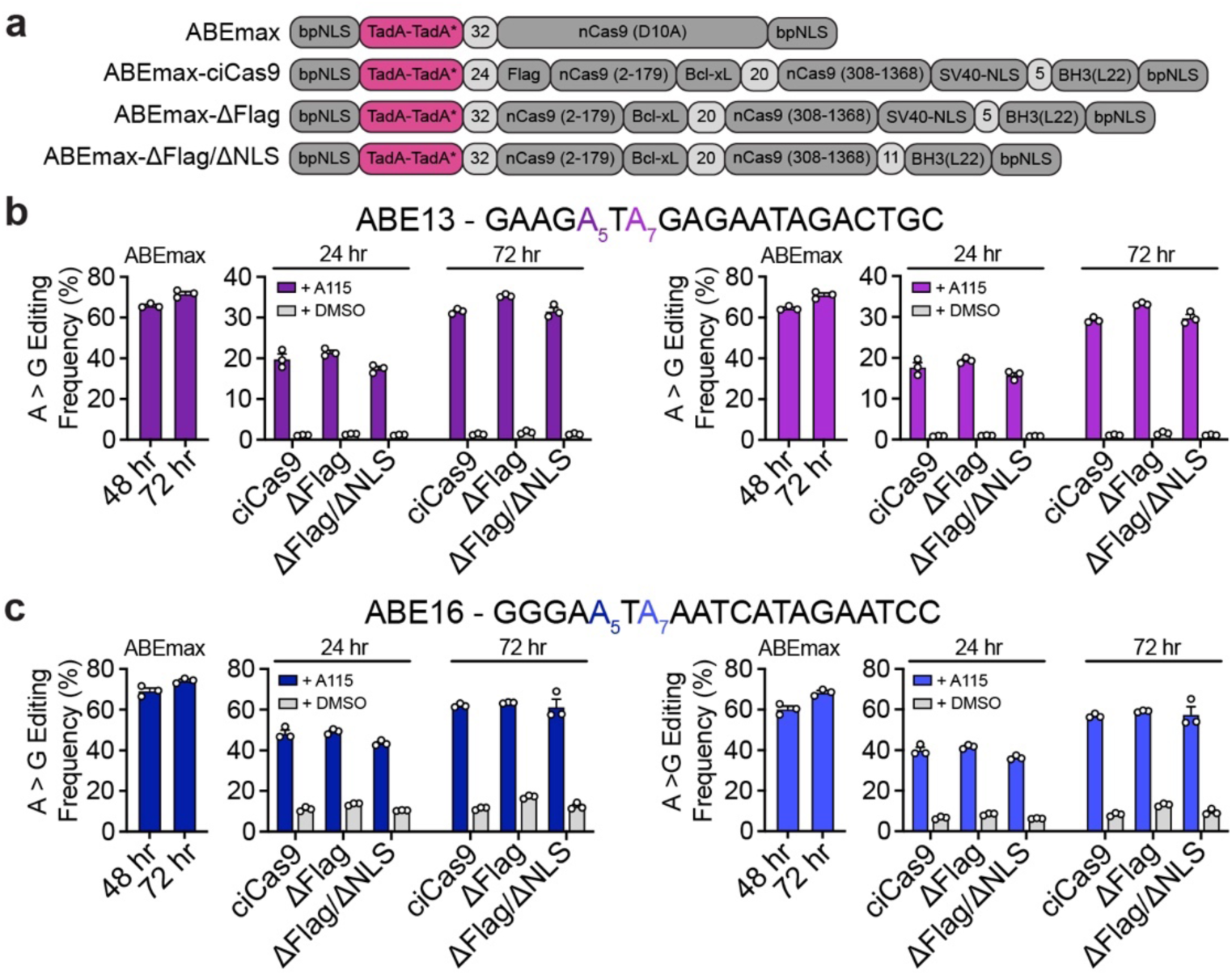
Chemically-controlled adenine base editors without codon optimization. **a)** Schematic of the domain arrangements in the unmodified ABEmax base editor and the chemically-controlled ciABEmax base editor without codon optimization and using the ciCas9(L22) variant. 3 different versions of ciCas9 were used, ciCas9(L22), ciCas9(L22) without a Flag-tag (ΔFlag), and ciCas9(L22) without a Flag-tag and additional SV40-NLS (ΔFlag/ΔNLS). **b-c)** A-to-G editing frequency with ABEmax and ABEmax-ciCas9 at the ABE13 **(b)** and ABE16 **(c)** target sites. ABEmax editing was measured at 48 and 72 hr after co-transfection of ABEmax and sgRNA. ABEmax-ciCas9 editing was measured at 24 and 72 hr after 1 μM A115 addition. A-to-G editing is shown at the 2 nucleotides in each target site with highest editing frequency with the Cas9 version of the ABEmax base editor. The 2 different nucleotides are indicated by color in the target sequence. Bars show mean editing frequency ± SEM of 3 cell culture replicates with white circles showing individual replicates.

**Supplemental Figure 7.**
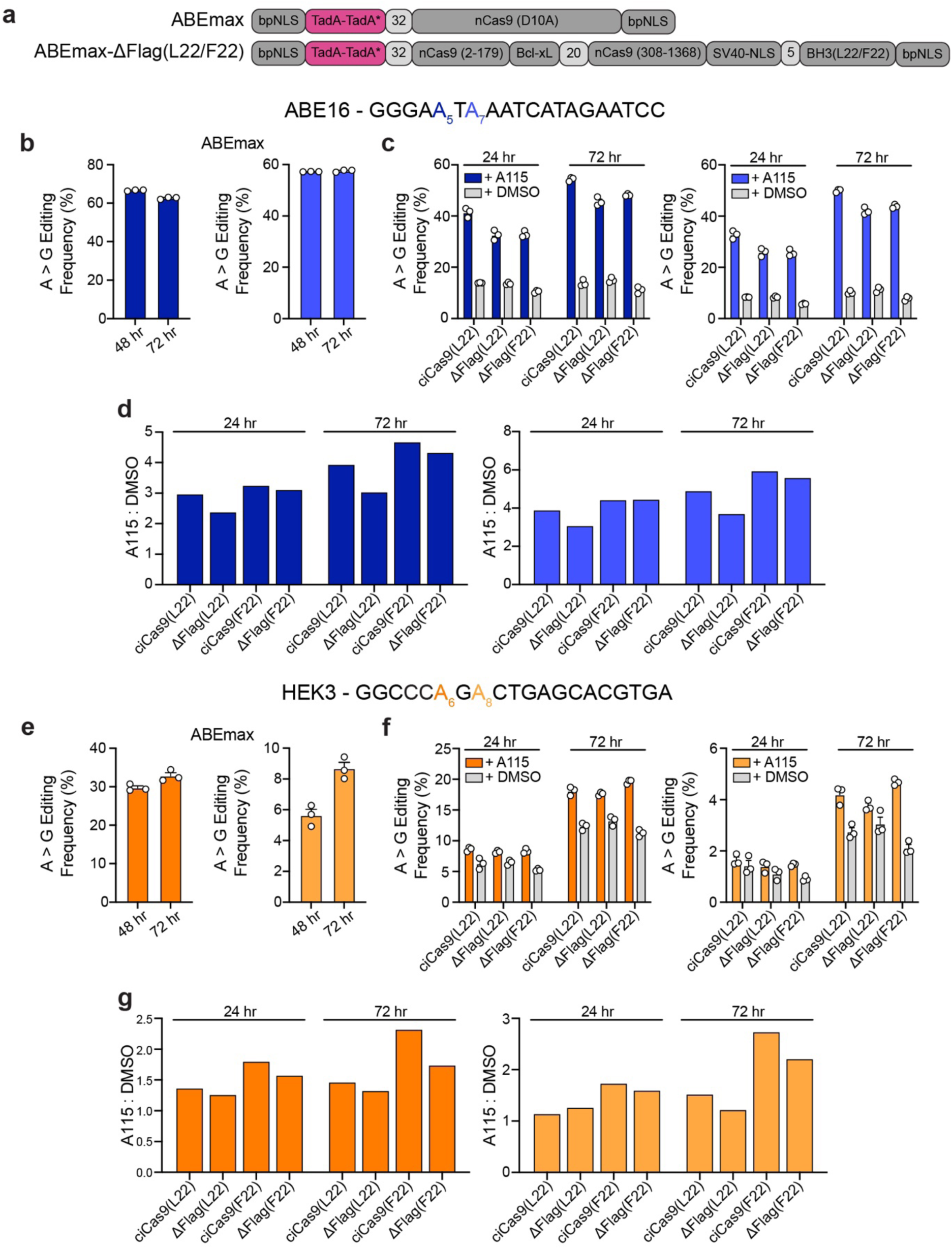
Additional constructs of codon optimized chemically-controlled ABEmax editors. **a)** Schematic of domains in the unmodified ABEmax base editor and additional constructs of the codon optimized ABEmax-ciCas9 base editors tested. 3 different versions of ciCas9 were **b,e)** A-to-G editing frequencies of the unmodified ABEmax base editor at the ABE16 **(b)** and HEK3 **(e)** target sites. **c,f)** A-to-G editing frequencies of the ABEmax-ciCas9 constructs at the ABE16 **(c)** and HEK3 **(f)** target sites. In **(b-c, e-f)** A-to-G editing is shown at the 2 nucleotides at each target site with highest editing frequency with ABEmax. The 2 different nucleotides are indicated by color in the target sequence. Editing by all ABEmax-ciCas9 constructs are quantified at 24 and 72 hr after 1 μM A115 or DMSO addition to HEK-293T cells. Bars show mean editing ± SEM of 3 cell culture replicates with white circles showing individual replicates. **d,g)** Ratio of the mean A-to-G editing frequency with 1 μM A115 to the mean A-to-G editing frequency with DMSO (A115:DMSO) for all tested ABEmax-ciCas9 base editors in Fig. 3 and Supplemental Fig. 7 at the ABE16 **(d)** and HEK3 **(g)** target sites. Bars show the ratios of editing at the 2 nucleotides at each target site with highest editing frequency with ABEmax. Editing frequencies used to calculate the ratio were measured at 24 and 72 hr after A115 addition to HEK-293T cells.

**Supplemental Figure 8.**
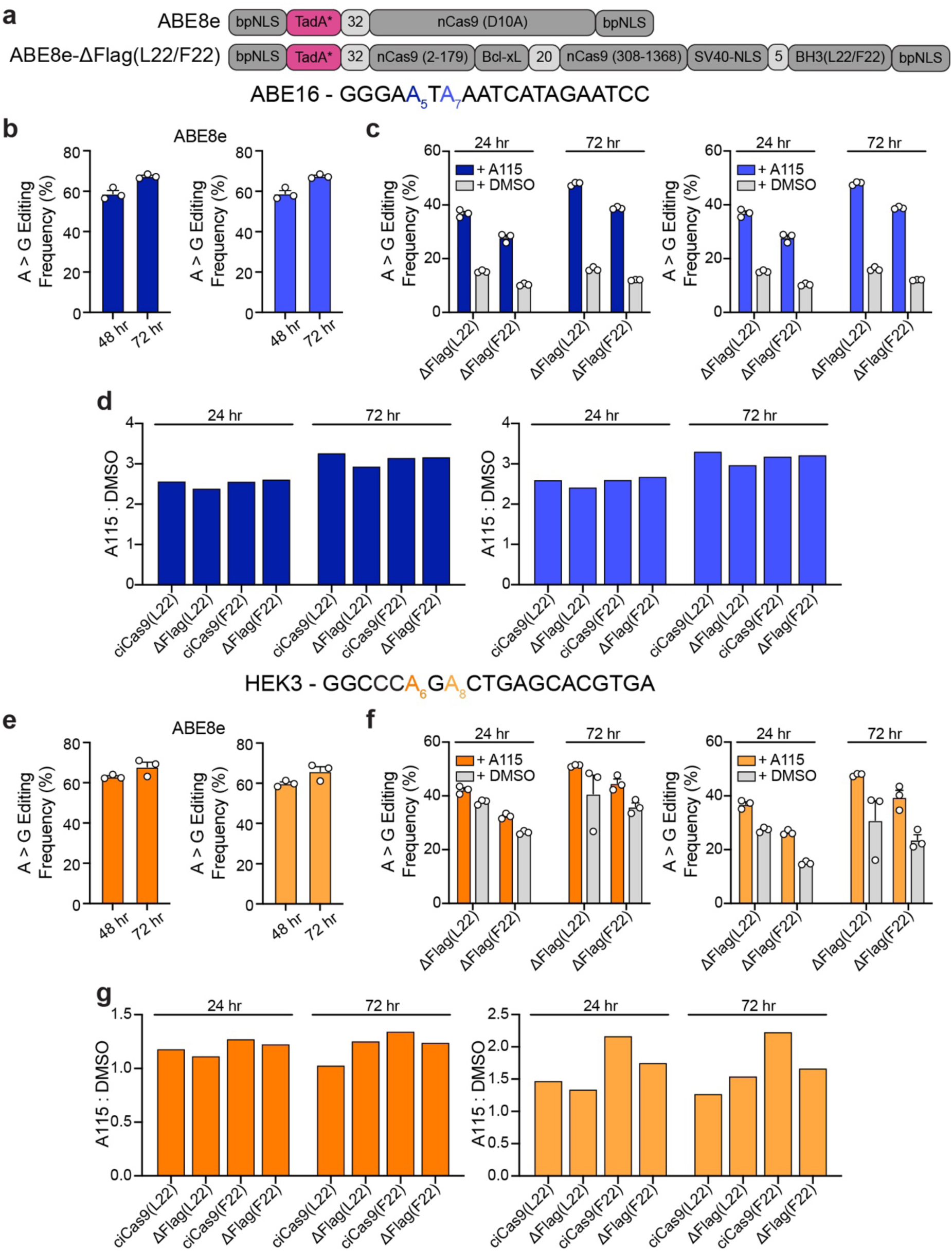
Additional constructs of codon optimized chemically-controlled ABE8e editors. **a)** Schematic of domains in the unmodified ABE8e base editor and additional constructs of the codon optimized ABE8e-ciCas9 base editors tested. 3 different versions of ciCas9 were additionally tested: ciCas9(L22), No-Flag-ciCas9(L22) (ΔFlag(L22)), and No-Flag-ciCas9(F22) **b,e)** A-to-G editing frequencies of the unmodified ABE8e base editor at the ABE16 **(b)** and HEK3 **(e)** target sites. **c,f)** A-to-G editing frequencies of the ABE8e-ciCas9 constructs at the ABE16 **(c)** and HEK3 **(f)** target sites. In **(b-c, e-f)** A-to-G editing is shown at the 2 nucleotides at each target site with highest editing frequency with ABE8e. The 2 different nucleotides are indicated by color in the target sequence. Editing by all ABE8e-ciCas9 constructs are quantified at 24 and 72 hr after 1 μM A115 or DMSO addition to HEK-293T cells. Bars show mean editing ± SEM of 3 cell culture replicates with white circles showing individual replicates. **d,g)** Ratio of the mean A-to-G editing frequency with 1 μM A115 to the mean A-to-G editing frequency with DMSO (A115:DMSO) for all tested ABE8e-ciCas9 base editors in Fig. 3 and Supplemental Fig. 8 at the ABE16 **(d)** and HEK3 **(g)** target sites. Bars show the ratios of editing at the 2 nucleotides at each target site with highest editing frequency with ABE8e. Editing frequencies used to calculate the ratio were measured at 24 and 72 hr after A115 addition to HEK-293T cells.

**Supplemental Figure 9.**
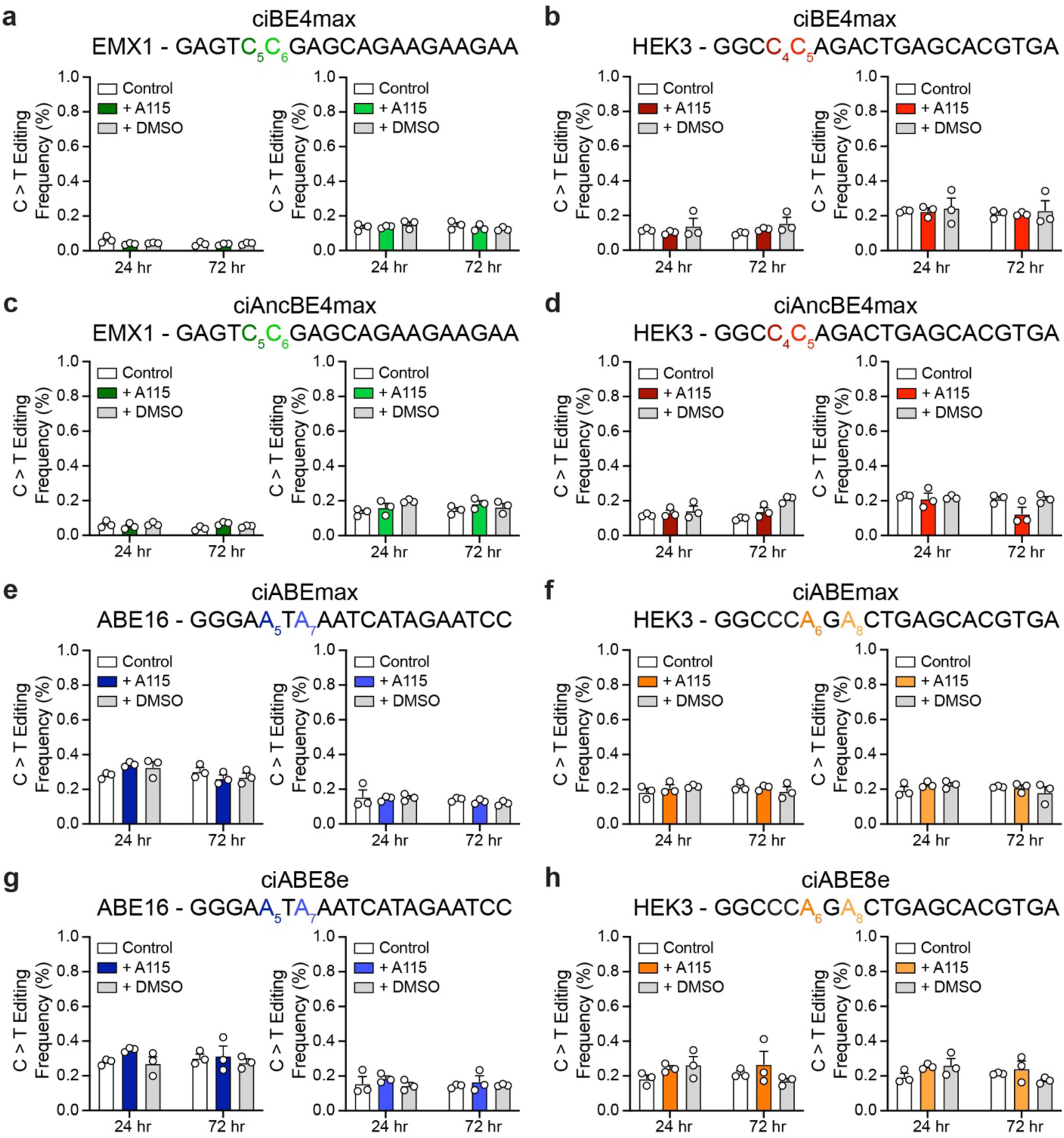
No sgRNA control for chemically-controlled base editors. **a-b)** ciBE4max base editing without sgRNA transfected at the EMX1 **(a)** and HEK3 **(b)** target sites. **c-d)** ciAncBE4max base editing without sgRNA transfected at the EMX1 **(c)** and HEK3 **(d)** target sites. **e-f)** ciABEmax base editing without sgRNA transfected at the ABE16 **(e)** and HEK3 **(f)** target sites. **g-h)** ciABE8e base editing without sgRNA transfected at the ABE16 **(g)** and HEK3 **(h)** target sites. More transfection control plasmid, pMAX-GFP, was used to replace the sgRNA plasmid in the cotransfection with base editor. C-to-T editing **(a-d)** and A-to-G editing **(e-h)** is shown at the 2 nucleotides at each target site with highest editing frequency with the Cas9 version of the base editor. Editing by all chemically-inducible base editor constructs are quantified at 24 and 72 hr after 1 μM A115 or DMSO addition to HEK-293T cells. Bars mean editing ± SEM of 3 cell culture replicates with white circles showing individual replicates.

**Supplemental Figure 10.**
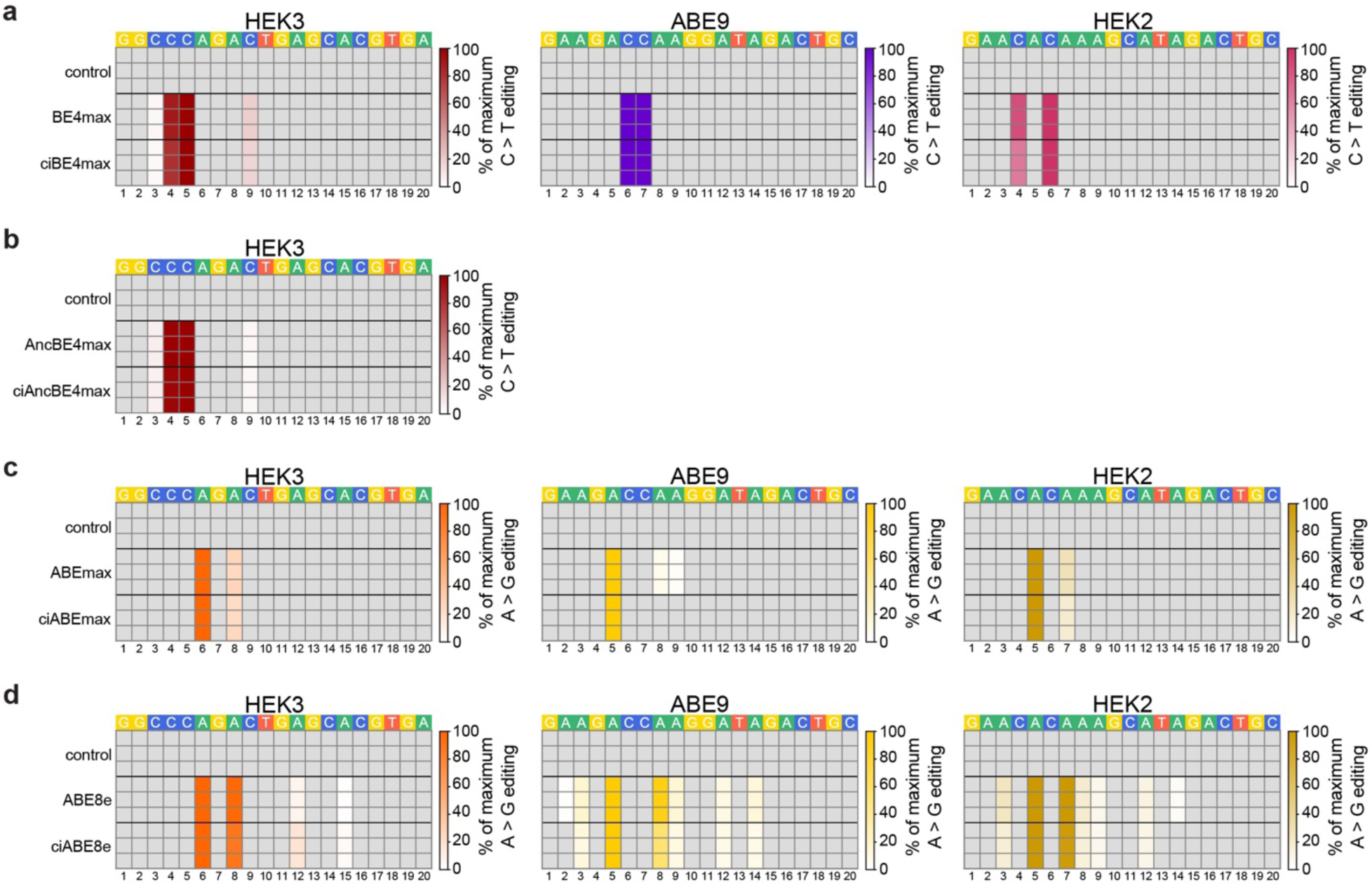
Heatmaps of base editing by chemically-controlled base editors compared to unmodified base editors. **a-b)** Heatmaps of BE4max, ciBE4max **(a)** and AncBE4max, ciAncBE4max **(b)** C-to-T base editing as a percentage of the highest edited nucleotide for each editor throughout the entire indicated target sites. **c-d)** Heatmaps of ABEmax, ciABEmax **(c)** and ABE8e, ciABE8e **(d)** A-to-G base editing as a percentage of the highest edited nucleotide for each editor throughout the entire indicated target sites. Each row shows an individual cell culture replicate. Editing frequencies of the unmodified base editors were quantified at 72 hr after transfection for the HEK3 target site and 48 hr after transfection for the ABE9 and HEK2 target sites. Chemically-controlled base editing frequencies were quantified at 72 hr after 1 μM A115 addition to HEK-293T cells for the HEK3 target site and 24 hr after 1 μM A115 addition to HEK-293T cells for the ABE9 and HEK2 target sites. The control shows untransfected cells harvested at the same time as the chemically-controlled base editors. The numbers below the heatmaps show the position of the nucleotide from the most PAM-distal nucleotide.

**Supplemental Figure 11.**
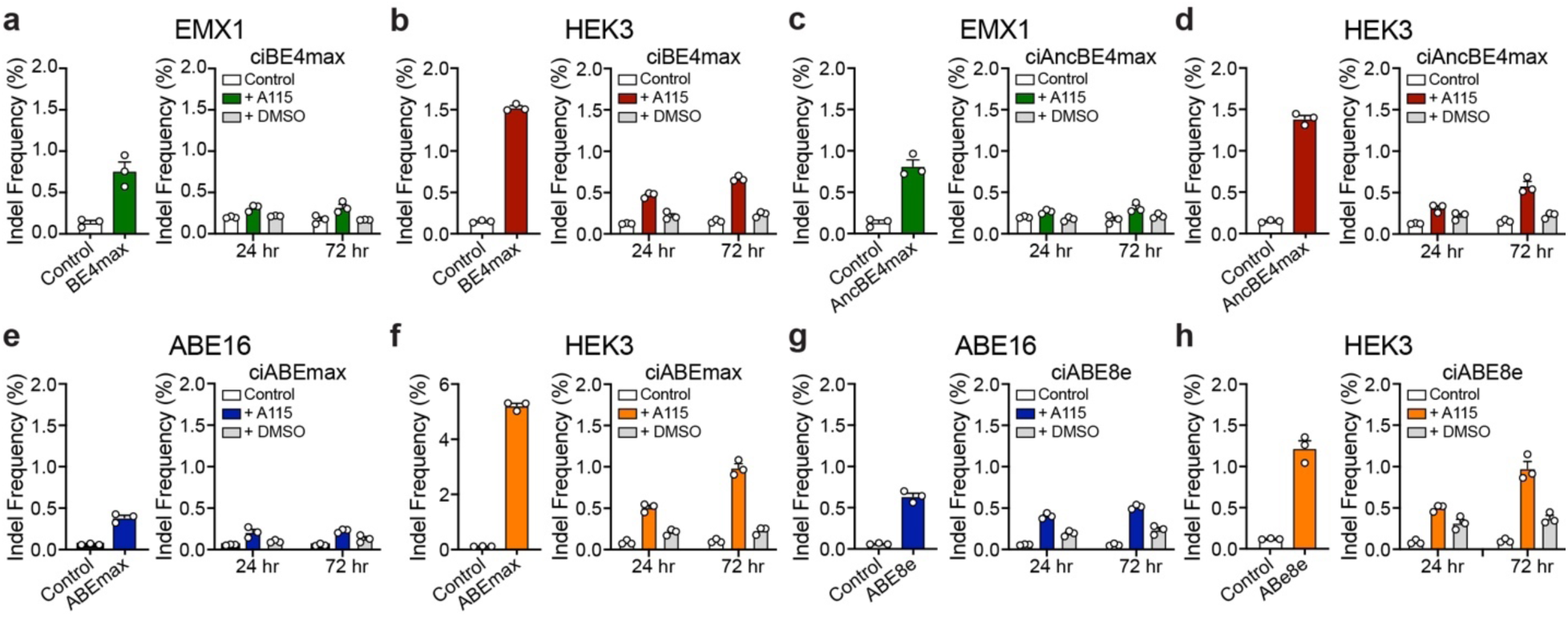
Indel formation by chemically-controlled base editors. **a-b)** BE4max (left) and ciBE4max (right) induced indel formation at the EMX1 **(a)** and HEK3 **(b)** target sites. **c-d)** AncBE4max (left) and ciAncBE4max (right) induced indel formation at the EMX1 **(c)** and HEK3 **(d)** target sites. **e-f)** ABEmax (left) and ciABEmax (right) induced indel formation at the ABE16 **(e)** and HEK3 **(f)** target sites. **g-h)** ABE8e (left) and ciABE8e (right) induced indel formation at the ABE16 **(g)** and HEK3 **(h)** target sites. Control samples were untransfected HEK-293T cells harvested at the same time as transfected cells. Editing by all unmodified base editors is quantified at 72 hr after transfection. Editing by all chemically-controlled base editors are quantified at 24 and 72 hr after 1 μM A115 or DMSO addition to HEK-293T cells. Bars show mean editing ± SEM of 3 cell culture replicates with white circles showing individual replicates.

**Supplemental Figure 12.**
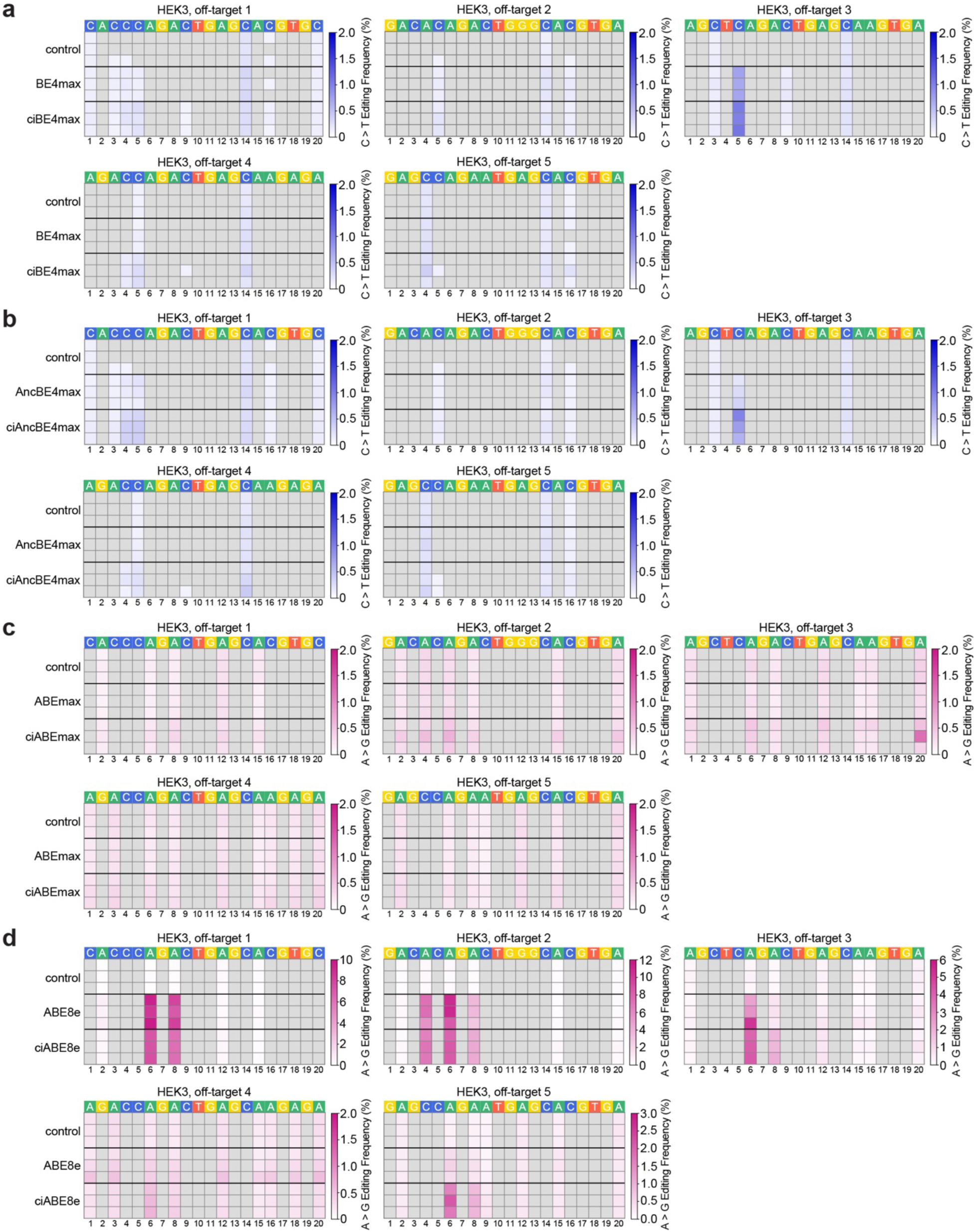
Off-target base editing by chemically-controlled base editors. Heatmaps of off-target base editing by ciBE4max **(a)**, ciAncBE4max **(b)**, ciABEmax **(c)**, and ciABE8e **(d)** with untransfected control and unmodified base editors. Each row shows an individual cell culture replicate. Unmodified base editor editing frequencies were quantified at 72 hr after transfection and chemically-controlled base editor editing frequencies were quantified at 72 hr after 1 μM A115 addition to HEK-293T cells. Untransfected control cells were harvested at the same time as chemically-controlled base editing cells. C-to-T and A-to-G base editing frequencies have been filtered to only include C or A nucleotides in the target site where >0.1% of base conversion is observed. The numbers below the heatmaps show the position of the nucleotide from the most PAM-distal nucleotide.

**Supplemental Figure 13.**
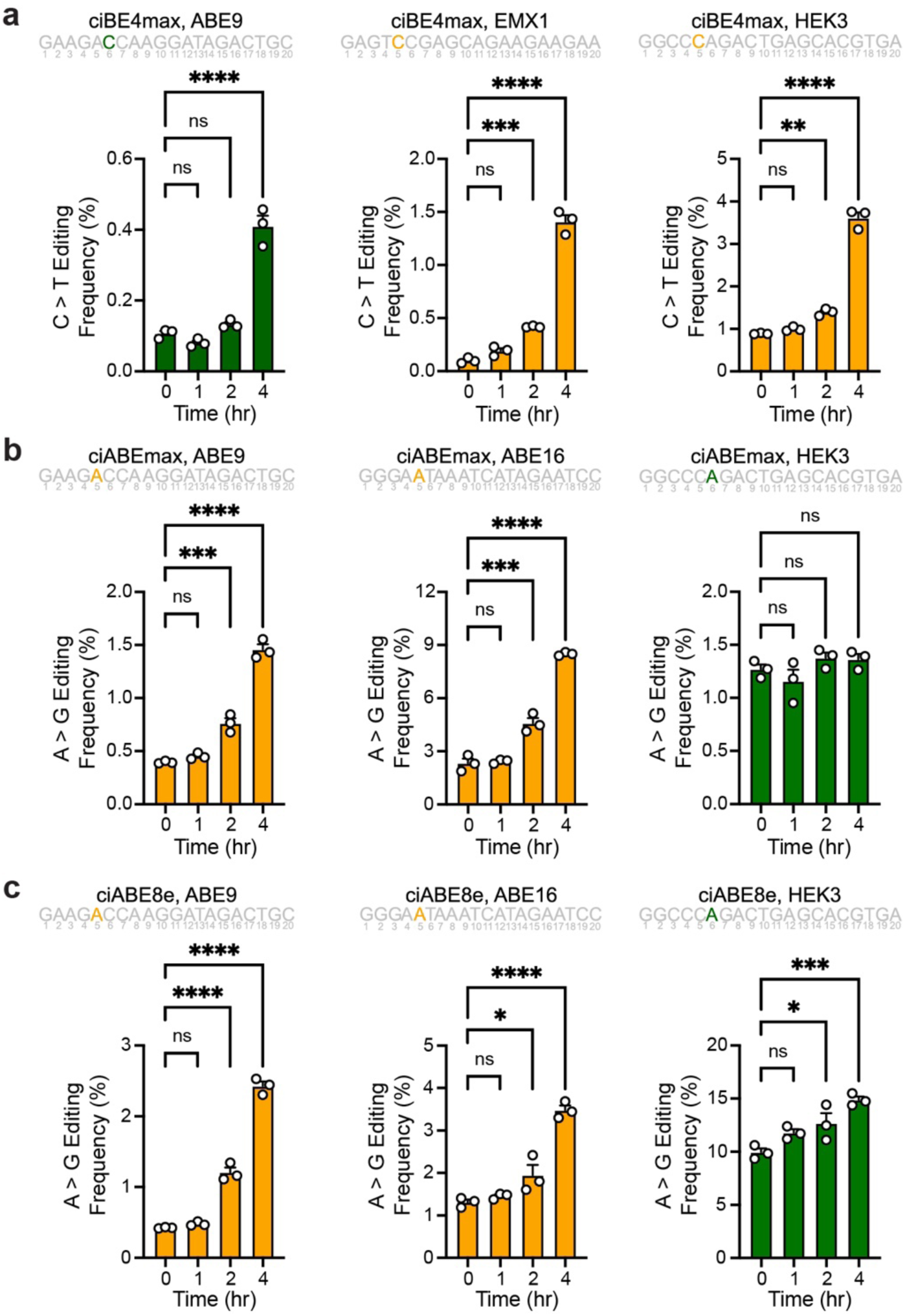
Early time points in time courses of base editing with the chemically-controlled base editors. Early time courses of chemically-controlled base editing using ciBE4max **(a)**, ciABEmax **(b)**, and ciABE8e **(c)** activated using 1 μM A115 at the indicated target sites. Time courses shown for the nucleotide colored in the target sequences shown. Numbers underneath the target sequence show the position of the nucleotide from the most PAM-distal nucleotide. Bars show mean editing ± SEM of 3 cell culture replicates with white circles showing individual replicates. Significance of editing at different time points were compared to editing frequency at 0 hr using a One-way ANOVA, statistical values shown in Supplemental Table 2.

**Supplemental Figure 14.**
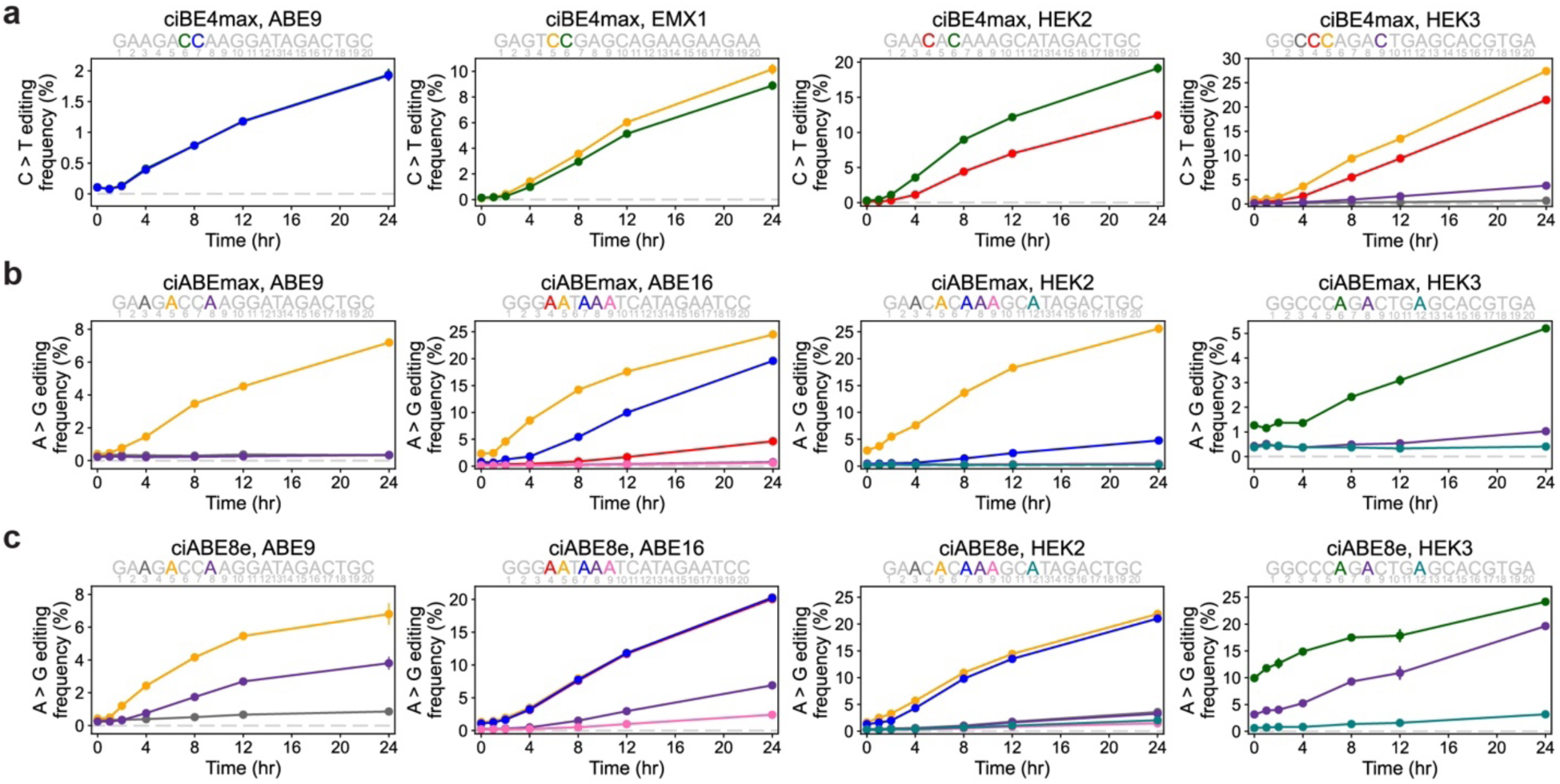
Time courses of base editing with the chemically-controlled base editors. **a)** Time course of chemically-controlled cytidine base editing by ciBE4max at the ABE9, EMX1, HEK2, and HEK3 target sites. ciBE4max was activated with 1 μM A115. Cells were harvested and editing was quantified at specified time points after activation. Colors of lines represent the corresponding nucleotide within the target site. Numbers underneath the target sequence show the position of the nucleotide from the most PAM-distal nucleotide. **b, c)** Time course of chemically-controlled adenine base editing by ciABEmax **(b)** and ciABE8e **(c)** at the ABE9, ABE16, HEK2, and HEK3 target sites. ciABEmax and ciABE8e were activated with 1 μM A115. Cells were harvested and editing was quantified at specified time points after activation. Colors of lines represent the corresponding nucleotide within the target site. Numbers underneath the target sequence show the position of the nucleotide from the most PAM-distal nucleotide. Data represented as mean editing ± SEM of 3 cell culture replicates. Time courses shown for all nucleotides where base editing frequency was greater than 0.5% at 24 hr after A115 addition.

**Supplemental Figures 15.**
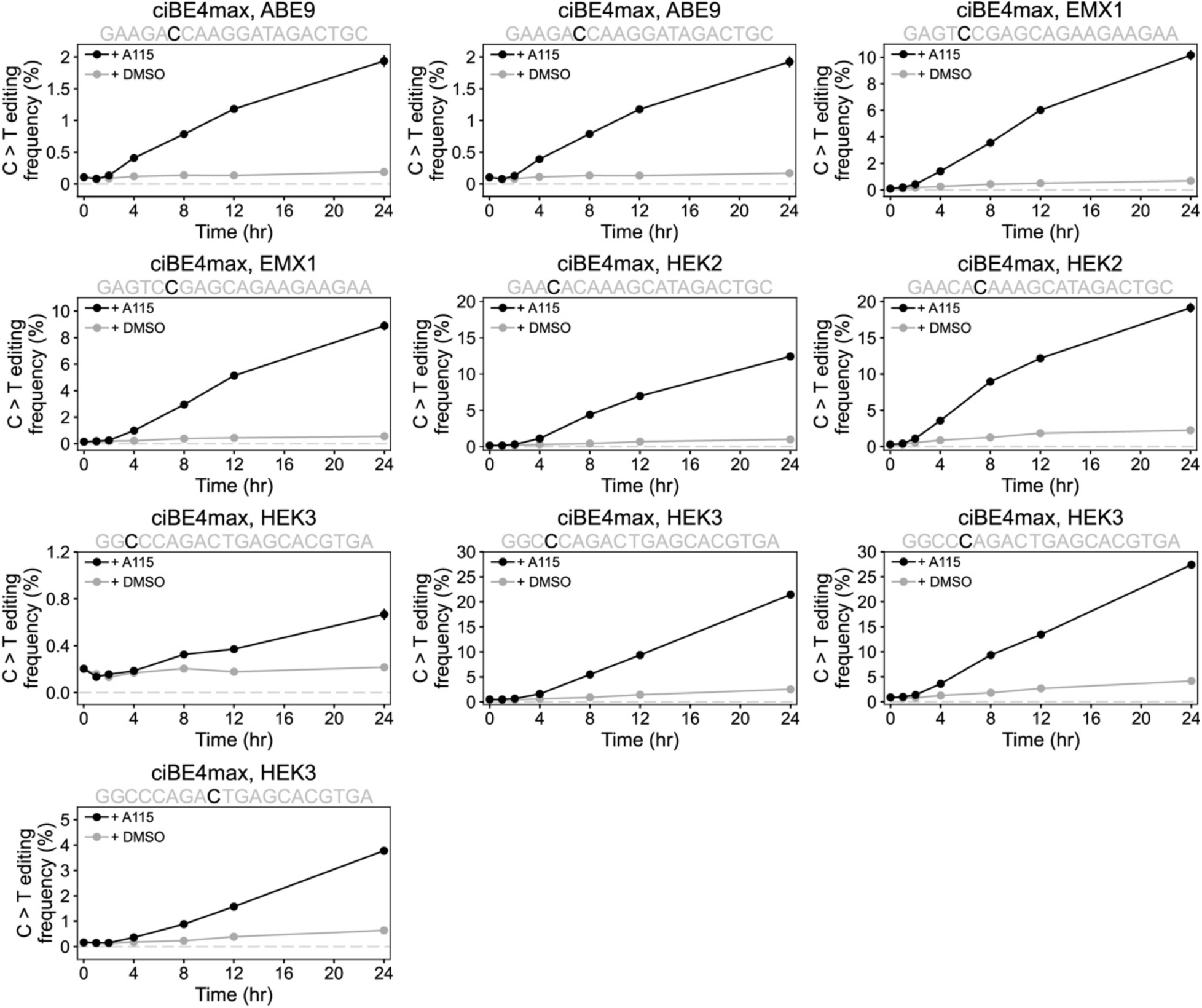
Time courses of ciBE4max base editing at individual nucleotides with A115 or DMSO. Time courses of ciBE4max C-to-T base editing. ciBE4max was activated with 1 μM A115 or DMSO. Cells were harvested and editing was quantified at the specified time points after activation. Black lines and circles show ciBE4max editing with 1 μM A115, gray lines and circles show ciBE4max editing with DMSO. Data represented as mean editing frequency ± SEM of 3 cell culture replicates.

**Supplemental Figures 16.**
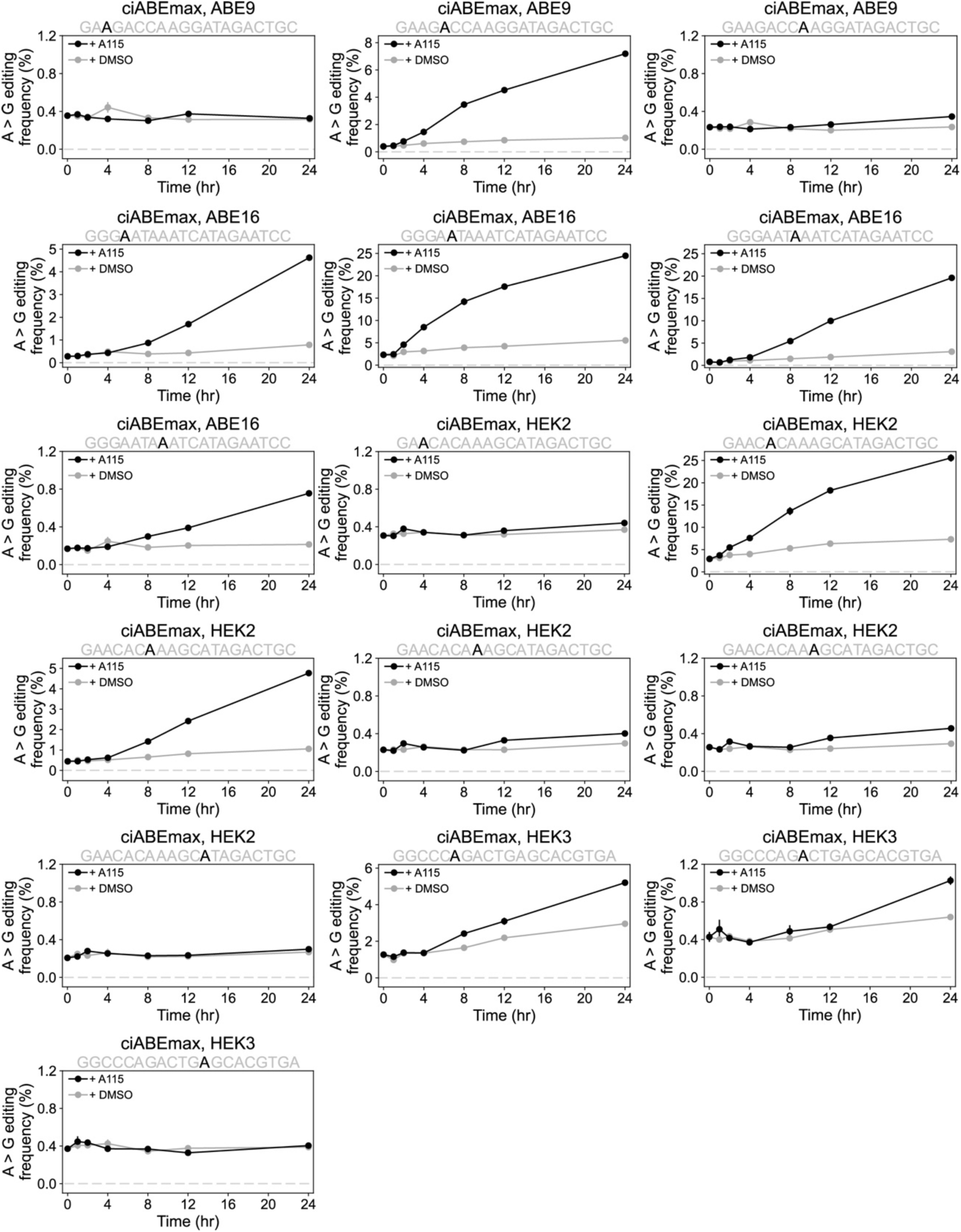
Time courses of ciABEmax base editing at individual nucleotides with A115 or DMSO. Time courses of ciABEmax C-to-T base editing. ciABEmax was activated with 1 μM A115 or DMSO. Cells were harvested and editing was quantified at the specified time points after activation. Black lines and circles show ciABEmax editing with 1 μM A115, gray lines and circles show ciABEmax editing with DMSO. Data represented as mean editing frequency ± SEM of 3 cell culture replicates.

**Supplemental Figures 17.**
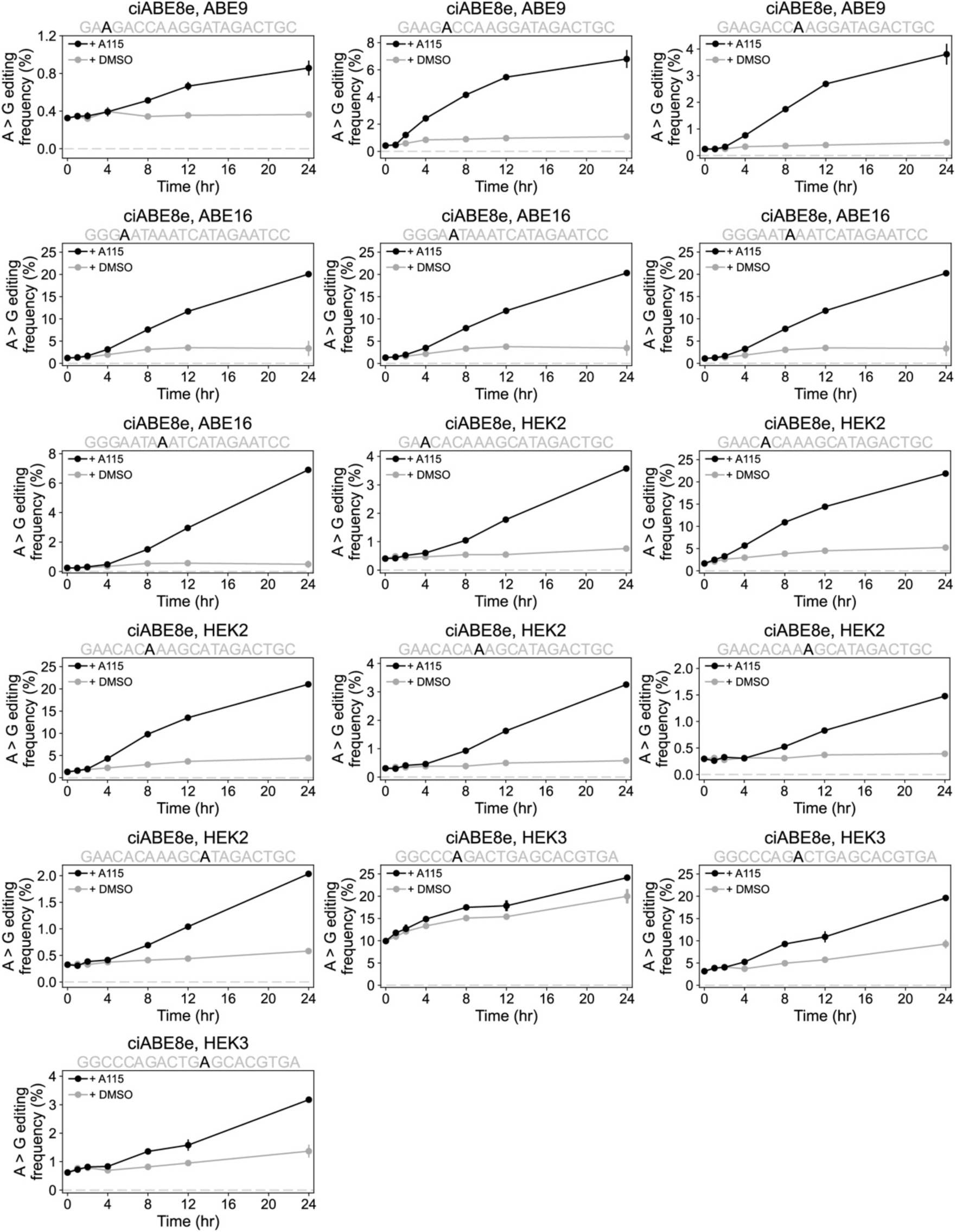
Time courses of ciABE8e base editing at individual nucleotides with A115 or DMSO. Time courses of ciABE8e C-to-T base editing. ciABE8e was activated with 1 μM A115 or DMSO. Cells were harvested and editing was quantified at the specified time points after activation. Black lines and circles show ciABE8e editing with 1 μM A115, gray lines and circles show ciABE8e editing with DMSO. Data represented as mean editing frequency ± SEM of 3 cell culture replicates.

**Supplemental Figure 18.**
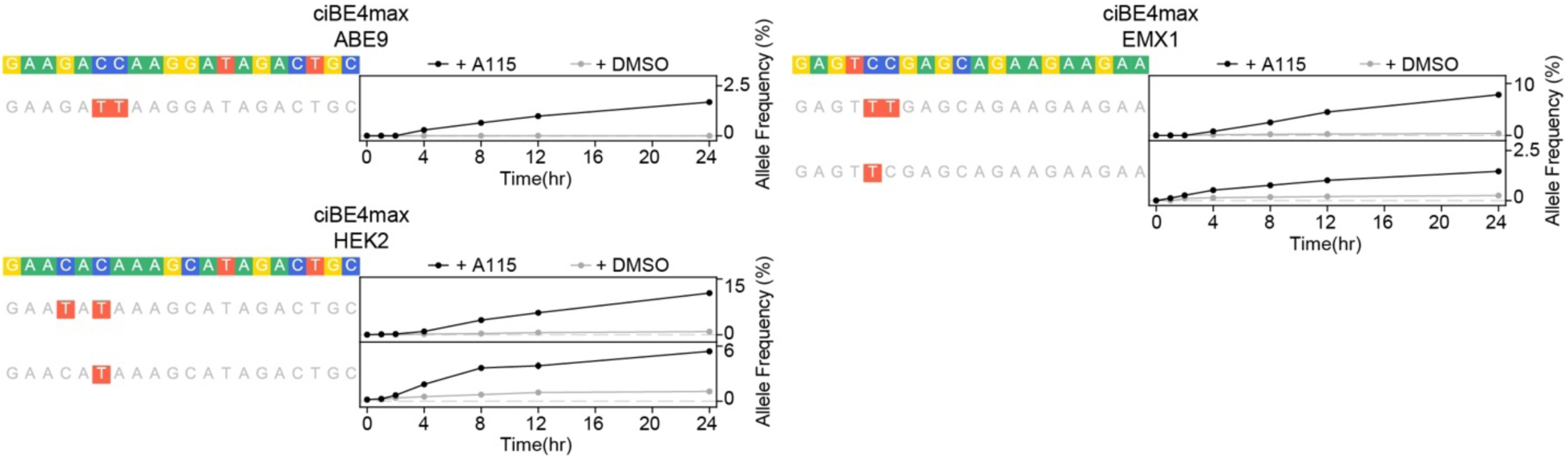
Time courses of ciBE4max base editing allele outcomes. Time course of allele formation by ciBE4max after activation with 1 μM A115 or DMSO. Black lines and circles show editing with 1 μM A115, gray lines and circles show editing with DMSO. Data represented as mean allele frequency ± SEM of 3 cell culture replicates.

**Supplemental Figure 19.**
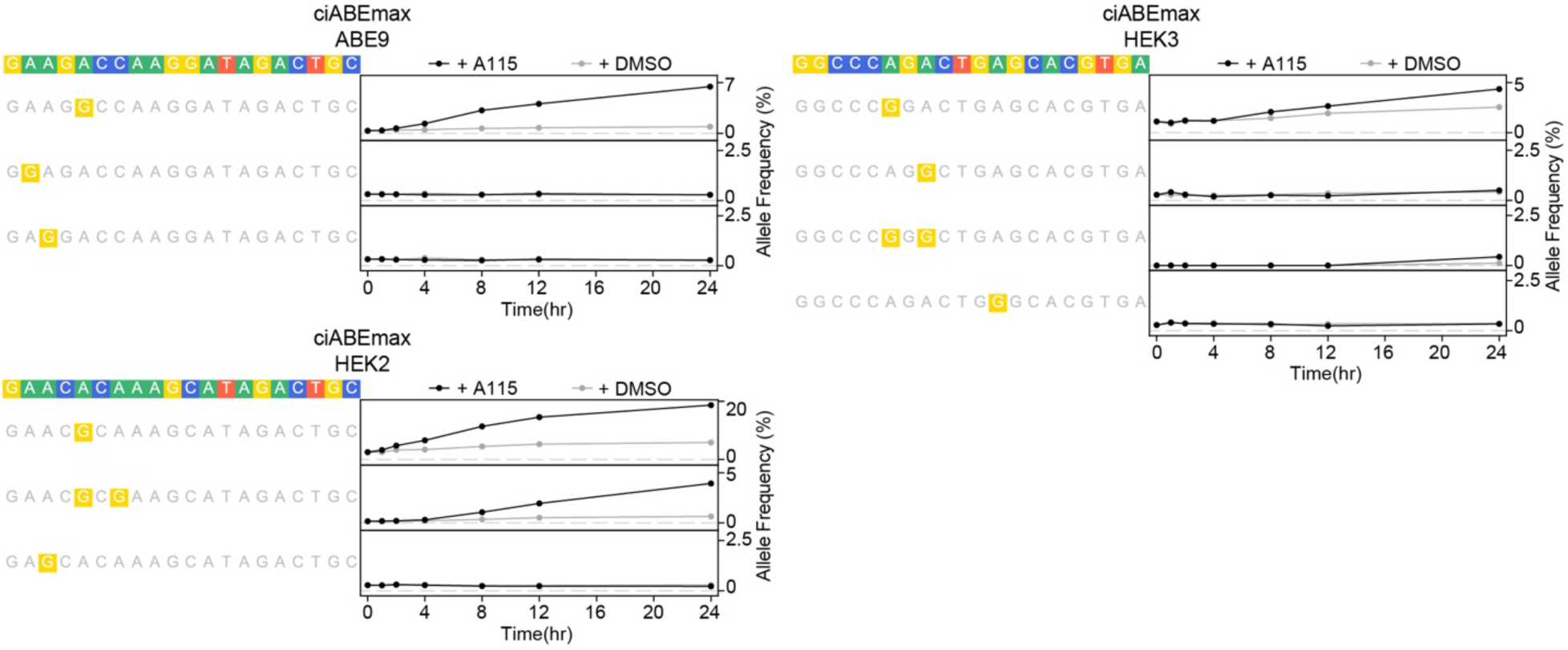
Time courses of ciABEmax base editing allele outcomes. Time course of allele formation by ciABEmax after activation with 1 μM A115 or DMSO. Black lines and circles show editing with 1 μM A115, gray lines and circles show editing with DMSO. Data represented as mean allele frequency ± SEM of 3 cell culture replicates.

**Supplemental Figure 20.**
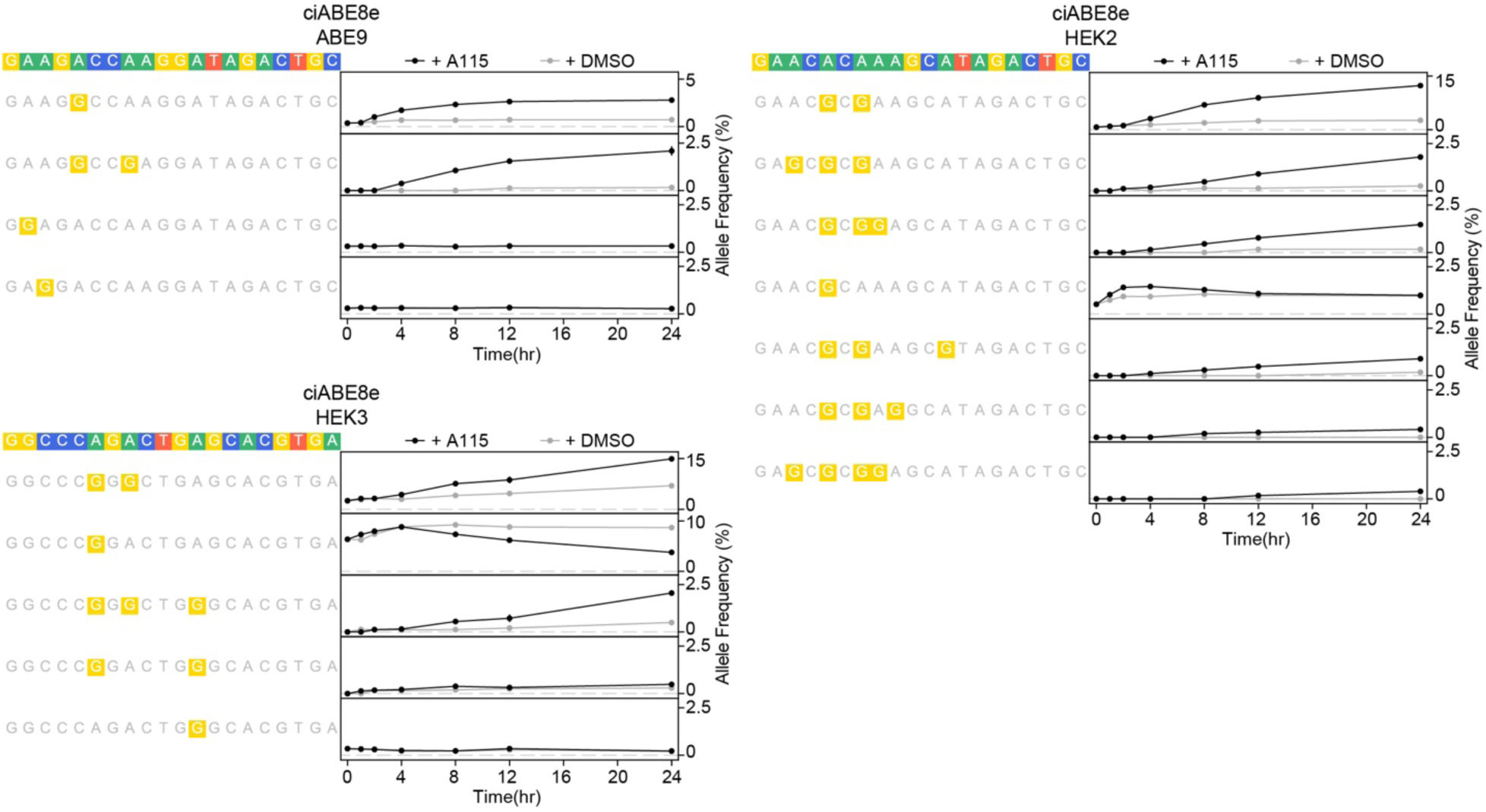
Time courses of ciABE8e base editing allele outcomes. Time course of allele formation by ciABE8e after activation with 1 μM A115 or DMSO. Black lines and circles show editing with 1 μM A115, gray lines and circles show editing with DMSO. Data represented as mean allele frequency ± SEM of 3 cell culture replicates.

**Supplemental Figure 21.**
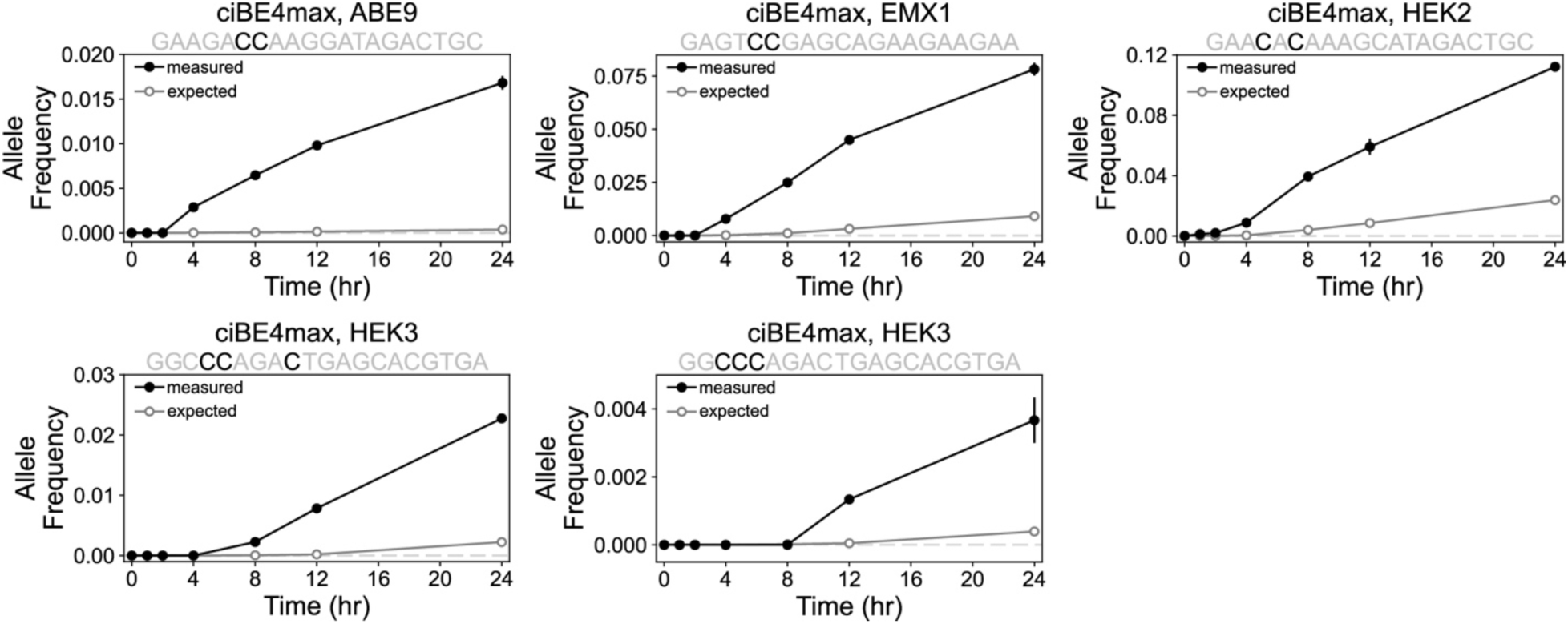
Time course of measured and expected allele frequencies by ciBE4max. Measured and expected allele frequencies over time created by ciBE4max that show a dependent model of base editing for multiply-edited alleles. Black lines and solid circles show measured allele frequencies, gray lines and open circles show expected allele frequencies. Measured data represented as mean editing frequency ± SEM of 3 cell culture replicates. Expected editing frequency represented as mean expected editing frequency ± relative error. Calculations for expected frequency and relative error described in the methods.

**Supplemental Figure 22.**
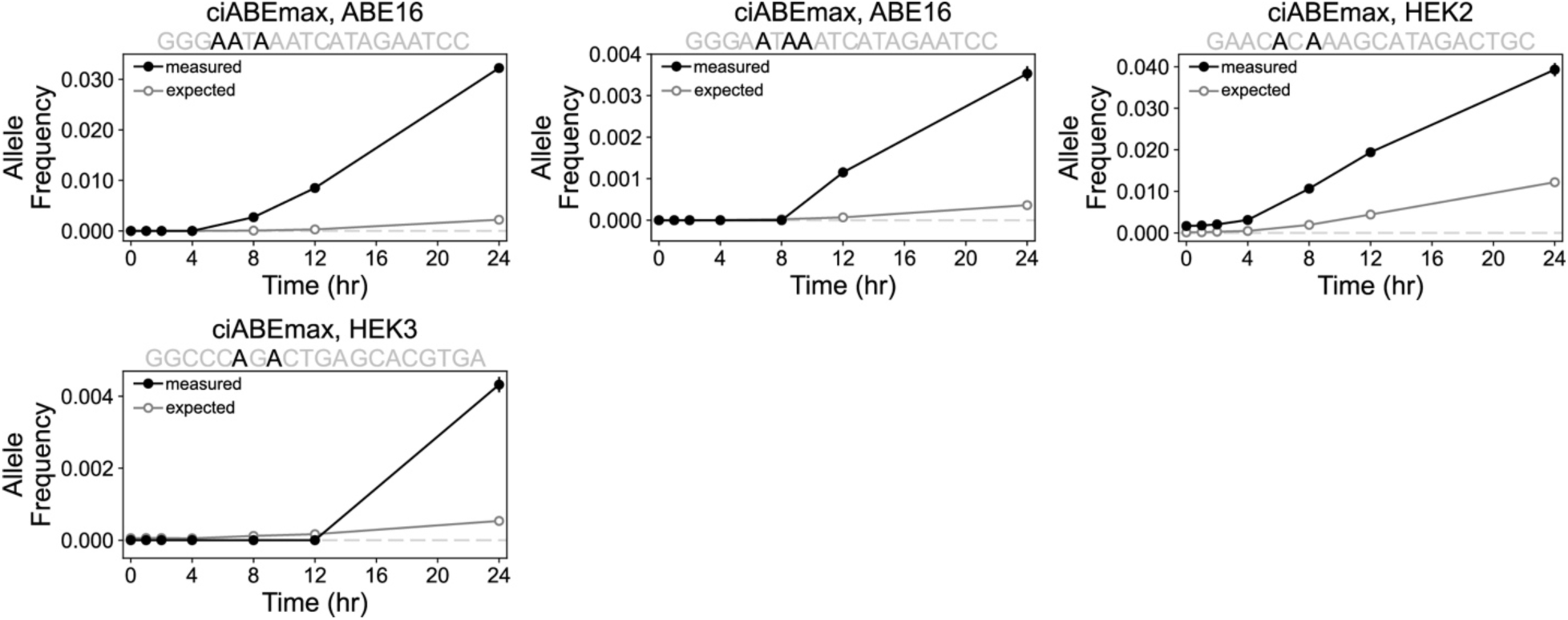
Time course of measured and expected allele frequencies by ciABEmax. Measured and expected allele frequencies over time created by ciABEmax that show a dependent model of base editing for multiply-edited alleles. Black lines and solid circles show measured allele frequencies, gray lines and open circles show expected allele frequencies. Measured data represented as mean editing frequency ± SEM of 3 cell culture replicates. Expected editing frequency represented as mean expected editing frequency ± relative error. Calculations for expected frequency and relative error described in Materials and Methods.

**Supplemental Figure 23.**
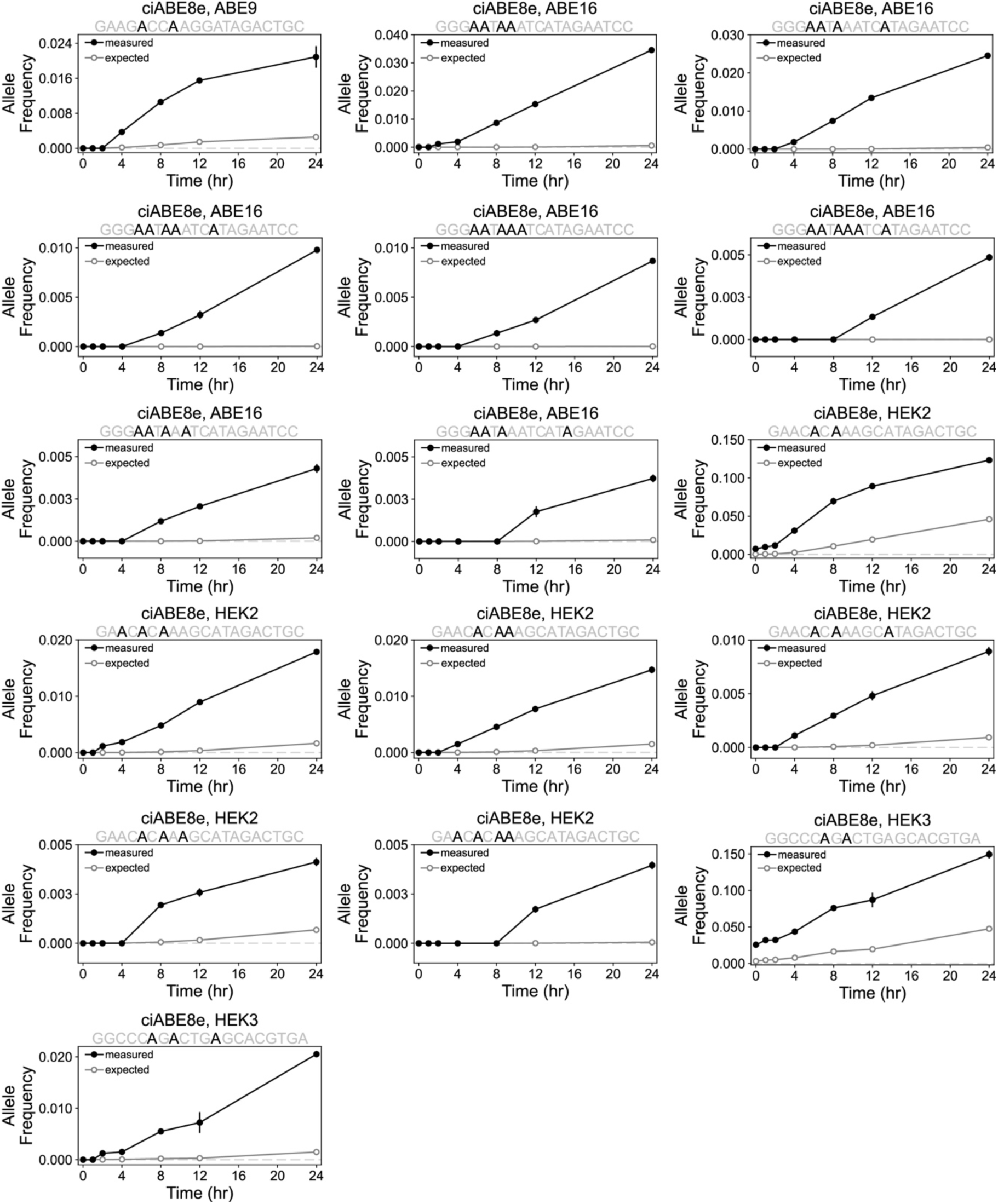
Time course of measured and expected allele frequencies by ciABE8e. Measured and expected allele frequencies over time created by ciABE8e that show a dependent model of base editing for multiply-edited alleles. Black lines and solid circles show measured allele frequencies, gray lines and open circles show expected allele frequencies. Measured data represented as mean editing frequency ± SEM of 3 cell culture replicates.Expected editing frequency represented as mean expected editing frequency ± relative error. Calculations for expected frequency and relative error described in Meterials and Methods.

**Supplemental Figure 24.**
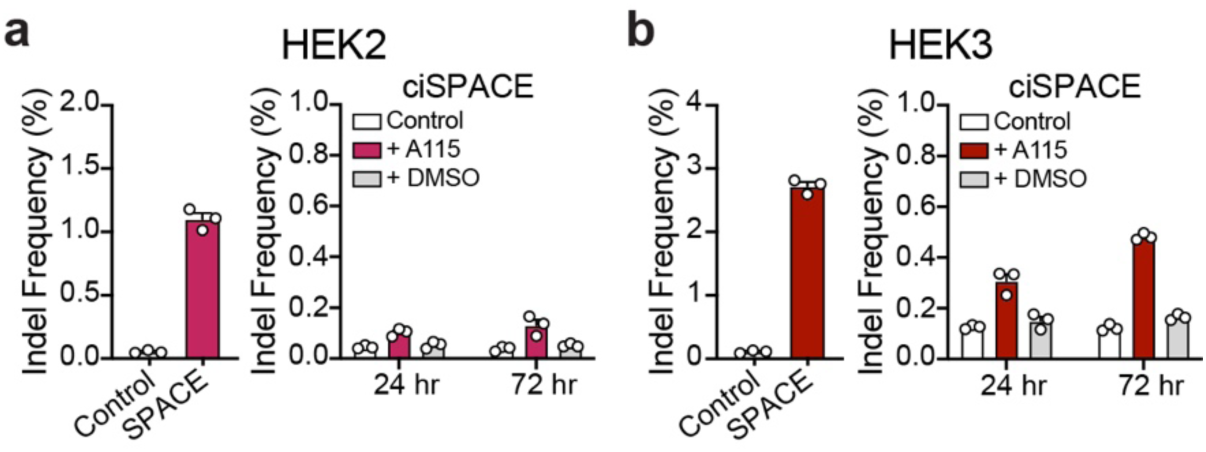
Indel formation by ciSPACE. SPACE (left) and ciSPACE (right) induced indel formation at the HEK2 **(a)** and HEK3 **(b)** target sites. Control samples were untransfected HEK-293T cells harvested at the same time as transfected cells. Editing by SPACE is quantified at 72 hr after transfection. Editing by ciSPACE is quantified at 24 and 72 hr after 1 μM A115 or DMSO addition to HEK-293T cells. Bars show mean editing ± SEM of 3 cell culture replicates with white circles showing individual replicates.

**Supplemental Figure 25.**
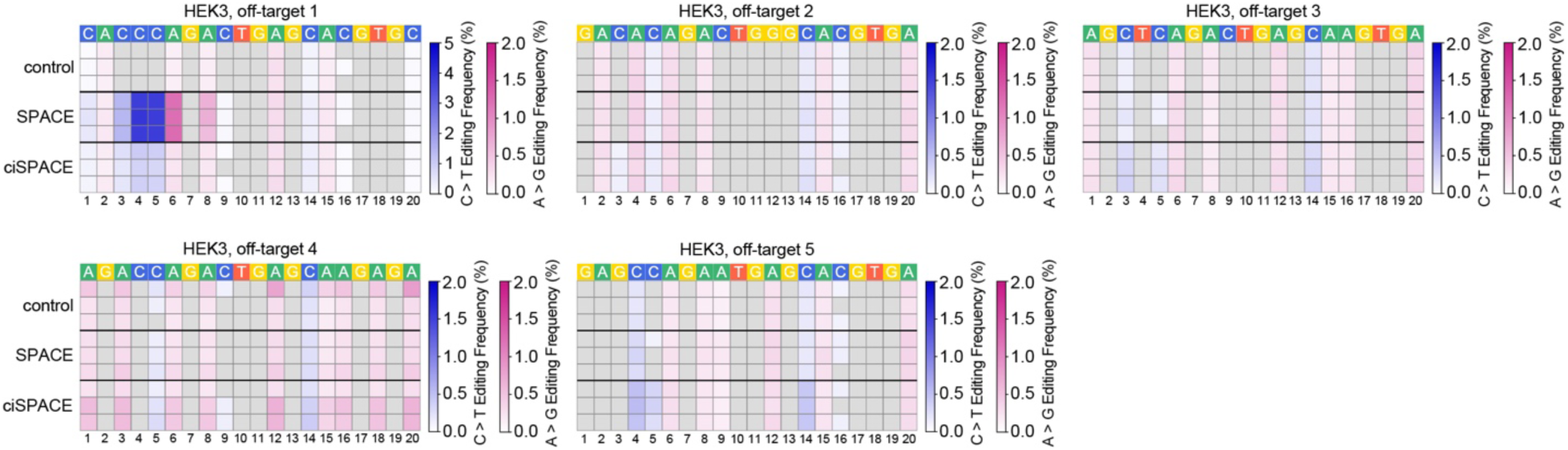
Off-target base editing by ciSPACE. Heatmaps of off-target base editing by ciSPACE with untransfected control and SPACE. Each row shows an individual cell culture replicate. SPACE editing frequencies were quantified at 72 hr after transfection and ciSPACE editing frequencies were quantified at 72 hr after 1 μM A115 addition to HEK-293T cells. Untransfected control cells were harvested at the same time as ciSPACE. C-to-T and A-to-G base editing frequencies have been filtered to only include C or A nucleotides in the target site where >0.1% of base conversion is observed. The numbers below the heatmaps show the position of the nucleotide from the most PAM-distal nucleotide.

**Supplemental Figure 26.**
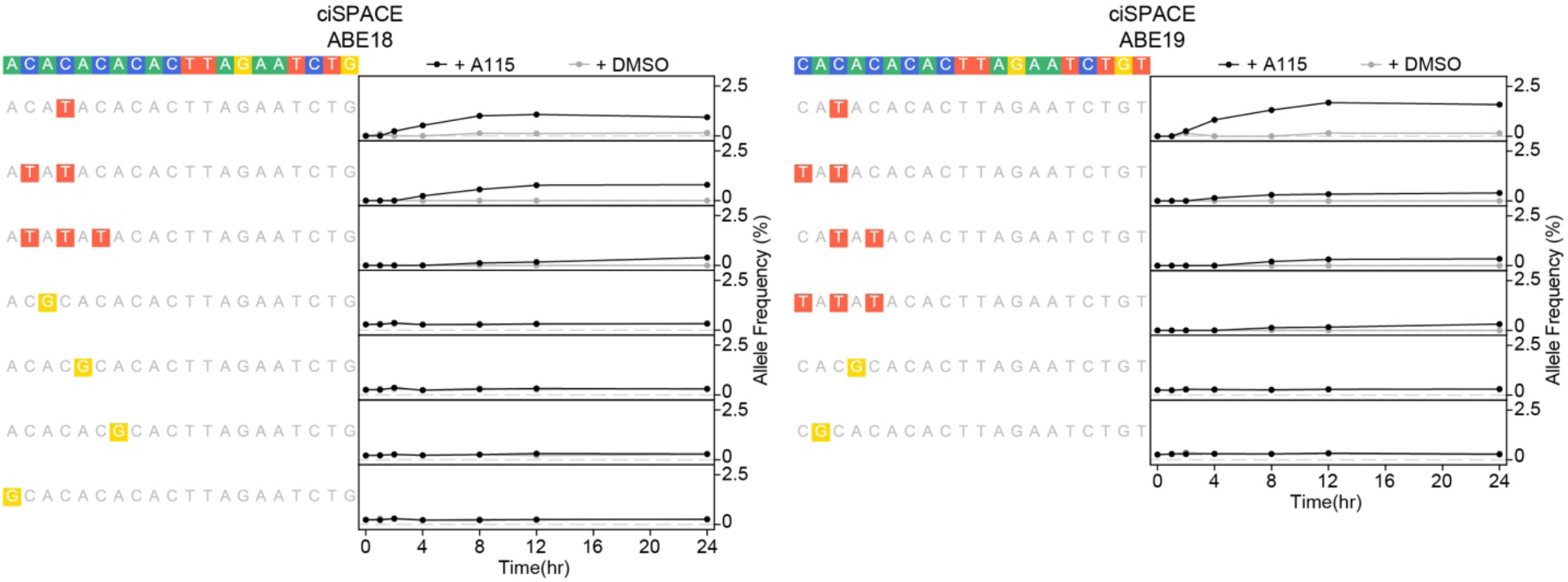
Allele frequency time courses by ciSPACE. Time course of allele formation by ciSPACE after activation with 1 μM A115 or DMSO. Black lines and circles show editing with 1 μM A115, gray lines and circles show editing with DMSO. Data represented as mean allele frequency ± SEM of 3 cell culture replicates.

**Supplemental Figure 27.**
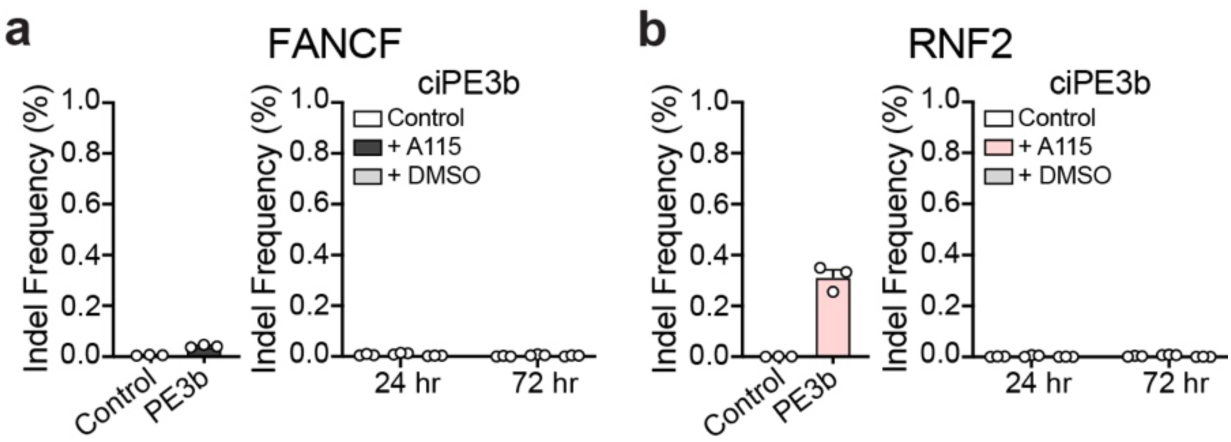
Indel formation by ciPE3b. PE3b (left) and ciPE3b (right) induced indel formation at the FANCF **(a)** and RNF2 **(b)** target sites corresponding to Fig. 6H. Control samples were untransfected HEK-293T cells harvested at the same time as transfected cells. Editing by PE3b is quantified at 72 hr after transfection. Editing by ciPE3b is quantified at 24 and 72 hr after 1 μM A115 or DMSO addition to HEK-293T cells. Bars show mean editing ± SEM of 3 cell culture replicates with white circles showing individual replicates.

**Supplemental Figure 28.**
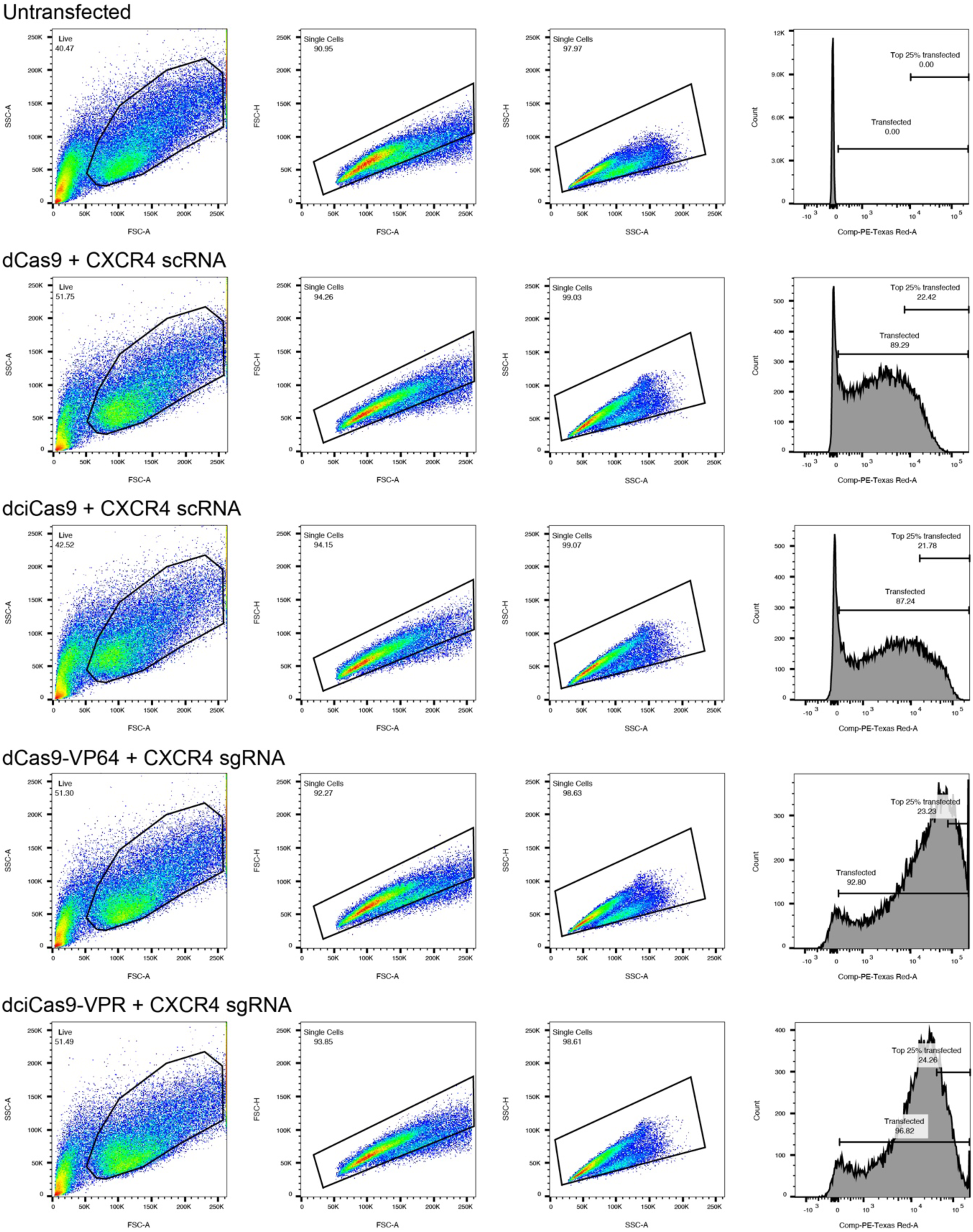
Example gating of transcriptional activation at the CXCR4 locus. Example gates set for determining CXCR4 transcriptional activation measured by median APC fluorescence of an anti-CXCR4 antibody, corresponding to Fig. 1C and Supplemental Fig. 1. HEK-293T cells were transfected with the indicated plasmids, outlined in the methods. Untransfected cells were stained with anti-CXCR4 antibody to determine background CXCR4 expression on HEK-293T cells.

**Supplemental Figure 29.**
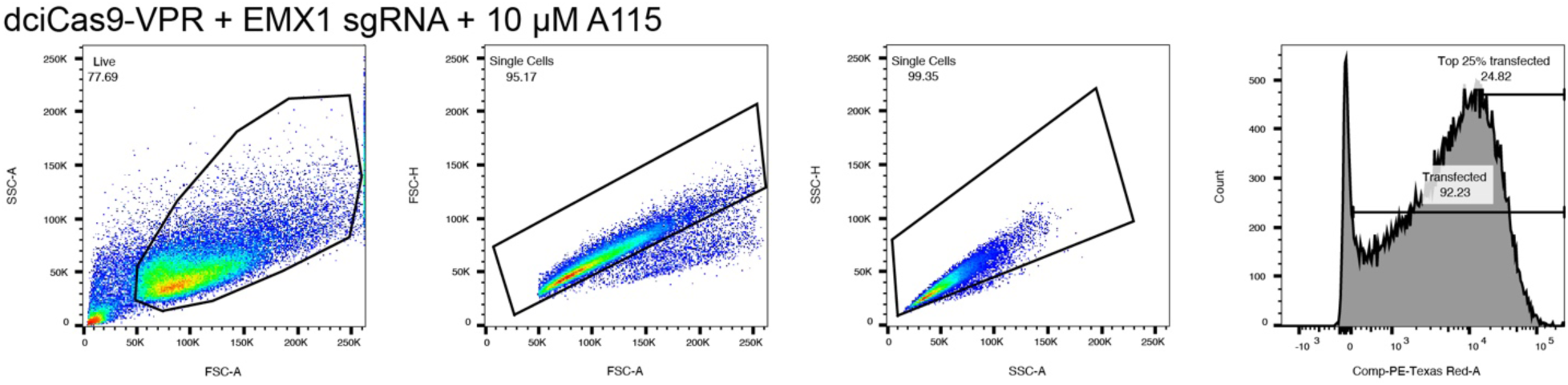
Example gating of transcriptional activation at the EMX1-EGFP synthetic reporter locus. Example gates set for determining EGFP transcriptional activation measured by EGFP fluorescence, corresponding to Fig. 1D. HEK-293 TREx FlpIn EMX1-EGFP cells were transfected with the indicated plasmids, outlined in the methods.

## SUPPLEMENTAL TABLES

**Supplemental Table 1.**
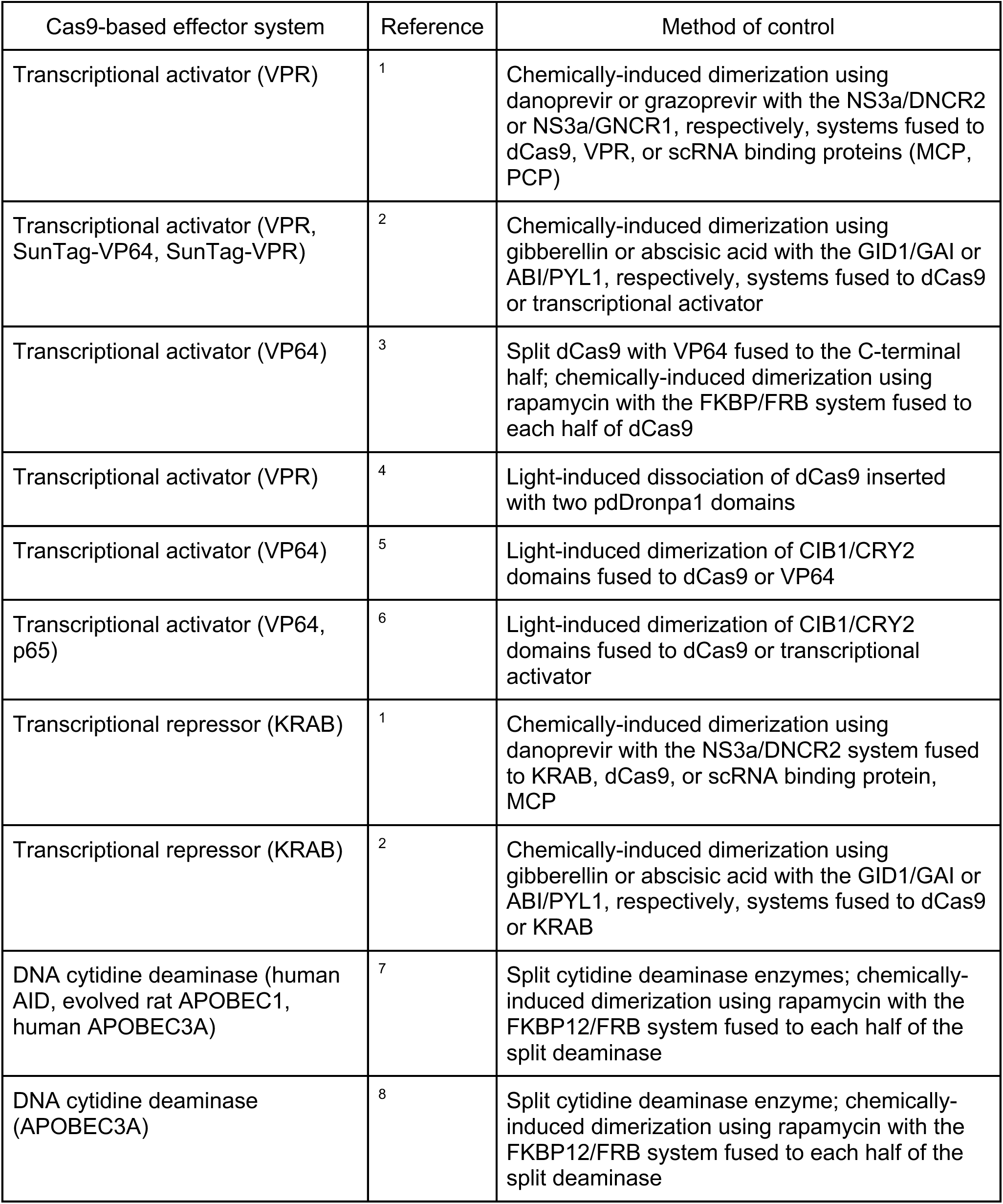
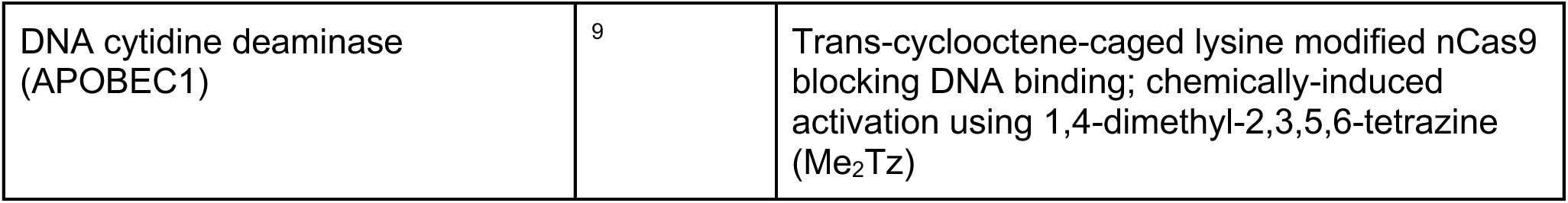
Summary of temporally-controlled Cas9-based effector systems.

| Cas9-based effector system | Reference | Method of control |
| --- | --- | --- |
| Transcriptional activator (VPR) | <sup>1</sup> | Chemically-induced dimerization using danoprevir or grazoprevir with the NS3a/DNCR2 or NS3a/GNCR1, respectively, systems fused to dCas9, VPR, or scRNA binding proteins (MCP, PCP) |
| Transcriptional activator (VPR, SunTag-VP64, SunTag-VPR) | <sup>2</sup> | Chemically-induced dimerization using gibberellin or abscisic acid with the GID1/GAI or ABI/PYL1, respectively, systems fused to dCas9 or transcriptional activator |
| Transcriptional activator (VP64) | <sup>3</sup> | Split dCas9 with VP64 fused to the C-terminal half; chemically-induced dimerization using rapamycin with the FKBP/FRB system fused to each half of dCas9 |
| Transcriptional activator (VPR) | <sup>4</sup> | Light-induced dissociation of dCas9 inserted with two pdDronpa1 domains |
| Transcriptional activator (VP64) | <sup>5</sup> | Light-induced dimerization of CIB1/CRY2 domains fused to dCas9 or VP64 |
| Transcriptional activator (VP64, p65) | <sup>6</sup> | Light-induced dimerization of CIB1/CRY2 domains fused to dCas9 or transcriptional activator |
| Transcriptional repressor (KRAB) | <sup>1</sup> | Chemically-induced dimerization using danoprevir with the NS3a/DNCR2 system fused to KRAB, dCas9, or scRNA binding protein, MCP |
| Transcriptional repressor (KRAB) | <sup>2</sup> | Chemically-induced dimerization using gibberellin or abscisic acid with the GID1/GAI or ABI/PYL1, respectively, systems fused to dCas9 or KRAB |
| DNA cytidine deaminase (human AID, evolved rat APOBEC1, human APOBEC3A) | <sup>7</sup> | Split cytidine deaminase enzymes; chemically-induced dimerization using rapamycin with the FKBP12/FRB system fused to each half of the split deaminase |
| DNA cytidine deaminase (APOBEC3A) | <sup>8</sup> | Split cytidine deaminase enzyme; chemically-induced dimerization using rapamycin with the FKBP12/FRB system fused to each half of the split deaminase |
| DNA cytidine deaminase (APOBEC1) | 9 | Trans-cyclooctene-caged lysine modified nCas9 blocking DNA binding; chemically-induced activation using 1,4-dimethyl-2,3,5,6-tetrazine (Me<sub>2</sub>Tz) |

**Supplemental Table 2.** One-way ANOVA results for comparison of early time points in chemically-controlled base editing.

| Base editor | Target site | Figure | One-way ANOVA comparison to 0 hr time point |  |  |  |  |  |
| --- | --- | --- | --- | --- | --- | --- | --- | --- |
|  |  |  | 1 hr |  | 2 hr |  | 4 hr |  |
|  |  |  | Mean difference | P-value | Mean difference | P-value | Mean difference | P-value |
| ciBE4max | ABE9 | Supp. Fig. 13a | 0.02648 | 0.5536 | -0.02574 | 0.5730 | -0.3029 | <0.0001 |
|  | EMX1 | Supp. Fig. 13a | -0.09143 | 0.2287 | -0.3205 | 0.0005 | -1.309 | <0.0001 |
|  | HEK2 | Fig. 4a | -0.1230 | 0.4713 | -0.8084 | <0.0001 | -3.274 | <0.0001 |
|  | HEK3 | Supp. Fig. 13a | -0.1094 | 0.6052 | -0.5002 | 0.0033 | -2.723 | <0.0001 |
| ciABEmax | ABE9 | Supp. Fig. 13b | -0.05634 | 0.5825 | -0.3672 | 0.0003 | -1.061 | <0.0001 |
|  | ABE16 | Supp. Fig. 13b | -0.1103 | 0.9623 | -2.260 | 0.0002 | -6.178 | <0.0001 |
|  | HEK2 | Fig. 4b | -0.7965 | 0.0476 | -2.565 | <0.0001 | -4.681 | <0.0001 |
|  | HEK3 | Supp. Fig. 13b | 0.1135 | 0.5533 | -0.1069 | 0.5943 | -0.09362 | 0.6781 |
| ciABE8e | ABE9 | Supp. Fig. 13c | -0.05563 | 0.7647 | -0.7806 | <0.0001 | -2.001 | <0.0001 |
|  | ABE16 | Supp.<br>Fig.<br>13c | -0.1471 | 0.8027 | -0.6336 | 0.0315 | -2.167 | <0.0001 |
|  | HEK2 | Fig. 4c | -0.8219 | 0.0121 | -1.609 | 0.0002 | -4.018 | <0.0001 |
|  | HEK3 | Supp.<br>Fig.<br>13c | -1.836 | 0.1199 | -2.740 | 0.0228 | -4.936 | 0.0007 |

**Supplemental Table 3. Chi-squared analysis results of significance in editing dependence.**

**Supplemental Table 4. List of all Cas9 and ciCas9 constructs with amino acid sequences, plasmid DNA sequences, and references.**

**Supplemental Table 5. gRNA sequences, primer sequences, and amplicon sequences.**

**Supplemental Table 6. Transcriptional activation median fluorescence values (for data presented in Figs. 1c-d and Supplemental Fig. 1).**

**Supplemental Table 7. DNA on-target sequencing data for non-codon optimized base editor experiments (for data presented in and Supplemental Figs. 3, 6).**

**Supplemental Table 8. DNA on-target sequencing data for codon optimized base editor experiments (for data presented in Figs. 2, 3, 6b-f, and Supplemental Figs. 4, 5, 7, 8, 9).**

**Supplemental Table 9. DNA base editor indel data (for data presented in Supplemental Figs. 11, 27).**

**Supplemental Table 10. DNA off-target sequencing data (for data presented in Supplemental Figs. 12, 25).**

**Supplemental Table 11. DNA sequencing data for time course experiments (for data presented in Fig. 4 and Supplemental Figs. 13-17).**

**Supplemental Table 12. Allele frequency data for time course experiments (for data presented in Figs. 5, 6g, and Supplemental Figs. 18-23, 26).**

**Supplemental Table 13. Calculated expected allele frequencies (for data presented in Figs. 5d-e and Supplemental Figs. 21-23).**

**Supplemental Table 14. Prime editing data and indel data (for data presented in Fig. 6h and Supplemental Fig. 27).**

